# Cathepsin K inhibitors promote osteoclast-osteoblast communication and engagement of osteogenesis

**DOI:** 10.1101/2025.03.04.641352

**Authors:** Preety Panwar, Jacob Bastholm Olesen, Jean-Marie Delaisse, Kent Søe, Dieter Brömme

## Abstract

Cathepsin K inhibitors are well known for their inhibitory activity against bone resorption but, in contrast with other bone resorption antagonists, were also reported to preserve bone formation in clinical trials. Here we show cathepsin K inhibitors favor the crosstalk between osteoclasts and osteoblasts and help engaging the osteogenic process required for proper bone remodeling. Therefore, we used a novel approach, co-culturing human osteoclasts and osteoblast lineage cells on bone slices and monitored through time-lapse their response to an active site (odanacatib) or an ectosteric (T06) cathepsin K inhibitor. Both inhibitors prevent the shift from pit to trench resorption mode and thus lead to a marked increase in pit-eroded surface lined with undigested collagen. Importantly, pit-eroded surfaces prove to receive significantly more and longer visits of osteoblast lineage cells. Furthermore, resorption achieved under CatK inhibition promotes osteoblast differentiation as shown by upregulation of alkaline phosphatase and type 1 collagen, and down regulation of RANKL. We propose a model where high cathepsin K activity levels lead to both aggressive bone resorption and compromised bone formation, and where low cathepsin K levels result in both slower resorption and faster initiation of formation. This model fits the current knowledge on the effect of collagen/collagenolysis on osteoclast activity and osteoblast chemotaxis. The combined effects of cathepsin K on resorption and formation render cathepsin K inhibitors unique tools to prevent bone loss. They stress the clinical interest of developing ectosteric inhibitors that may limit the side effects of active site inhibitors.

**LAY SUMMARY:** Small bone packages are continuously degraded by osteoclast cells and reconstructed by osteoblast cells. Too much degradation or too little reconstruction leads to bone loss and is currently treated with inhibitors of degradation or stimulators of reconstruction. There is usually little attention for the mechanism maintaining the balance between degradation and reconstruction. This mechanism involves proper communication between osteoclasts and osteoblasts. Here we show that cathepsin K inhibitors developed to inhibit degradation, also favor osteoclast-osteoblast communication, thereby allowing a faster preparation of degraded bone surfaces for new bone deposition. This highlights the unique clinical potential of cathepsin K inhibitors.

## INTRODUCTION

Bone remodeling is obligatory for the maintenance of the skeleton. It is achieved by remodeling units where osteoclasts (OCs) and osteoblast (OB) lineage cells work together to resorb and thereafter reconstruct the bone matrix. Close communication between OCs and OB lineage cells within this remodeling unit is important from the beginning of the remodeling cycle until the initiation of bone formation ^1^. OBs at an early stage of differentiation, such as reversal cells, activated bone lining cells and canopy cells exert a pro-resorptive effect on the OCs, as they produce factors like RANKL and MMP13 promoting OC-mediated bone resorption ^2–9^. In turn, OCs generate so-called coupling factors promoting OB cell expansion and differentiation on the bone surfaces they have eroded ^10,11^. While differentiating, these OB lineage cells (reversal cells) lose their pro-resorptive phenotype, become anti-resorptive ^6^, strongly alkaline phosphatase-positive (ALP^+^) ^12^, and start depositing collagen ^13^. Absence of OC-OB interactions results in lack of OB differentiation and bone reconstruction - leading to bone loss in diseases such as age-, menopause-, and glucocorticoid-induced osteoporosis ^1^. Thus, there is a need for understanding what determines OC-OB proximity, and what influences the duration and frequency of OC-OB physical interactions.

Here, we reasoned that changes in bone resorption and eroded surface characteristics may affect the approach of OB lineage cells to the OCs and their colonization of eroded surfaces. It has been repeatedly reported that OCs resorb bone according to two resorption modes, resulting in excavations appearing either as pits or trenches ^14–24^. The main determinant of the resorption mode was shown to be the level of cathepsin K (CatK) activity ^14^, which is driving the degradation of collagen by OCs. CatK inhibitors or low intrinsic CatK activity causes an accumulation of demineralized collagen, which stops the OC resorption machinery ^15^. The OC then migrates away soon after it started resorbing and leaves behind shallow pits lined with non-degraded demineralized collagen ^17,18,20^. In contrast, high CatK activity leads to long and deep resorption trenches devoid of demineralized collagen ^17,18,20^. Trenches are prevailing in pathophysiological situations of aggressive bone resorption ^25,26^. This mechanistic finding thus explains well the anti-resorptive efficacy of CatK inhibitors ^27^. Here, we hypothesize that these CatK inhibitor-induced changes in erosion may additionally affect “homing” of OB lineage cells onto the eroded surface^27^.

The active site-directed CatK inhibitors, odanacatib (ODN) and balicatib, showed inacceptable side effects in clinical trials,^27^ including fibrotic events in heart and skin ^28^. This might be related to an accumulation of TGF-ß in tissues when its degradation by CatK is inhibited by active site-directed inhibitors ^29^. Therefore, ectosteric inhibitors, which antagonize selectively the collagenolytic activity of CatK and still allow non-collagenous proteolytic activity of CatK including the degradation of TGF-ß, are now being developed ^17,18,30^. Interestingly, comparing both inhibitor types also sheds light on whether solely the collagenase activity of CatK is responsible for the anti-resorptive activity of CatK inhibitors. An ectosteric inhibitor, Tanshinone IIA sulfonate (T06), tested in a preclinical model inhibits bone resorption with comparable characteristics to ODN, but without interfering in non-collagen related activities of CatK^18,30^.

The present study is based on an OC-OB co-culture model that was designed to mimic the *in vivo* characteristics of reversal cells and has highlighted a pro-resorptive effect of OB lineage cells ^8^. In the current study, we combined this OC-OB co-culture model with time-lapse analysis to demonstrate the effect of ODN and T06 (i) on the frequency and duration of the associations of OB lineage cells with resorption events, (ii) on osteogenic markers, and (iii) on physical and microchemical characteristics of the excavations, which are likely to affect initiation of bone formation.

## MATERIALS AND METHODS

### OC preparation

CD14^+^ monocytes from blood of anonymous donors were differentiated into OCs as described ^31^. The use of anonymized buffy coats is in accordance with Danish legislation and all donors provided written consent for the use of surplus material. This procedure is approved by the Danish Ministry of Health and the Regional Ethics Committee of Southern Denmark. In brief, following Ficoll separation and purification with BD IMag™ Anti-Human CD14 magnetic particles (BD Biosciences, CA, USA), CD14^+^ cells were cultured in T75 flasks (5x10^6^ cells) in α-Minimum Essential Medium (αMEM; Invitrogen, Taastrup, Denmark) containing 10% fetal calf serum (FCS; Sigma-Aldrich, St Louis, MO) and 25 ng/ml recombinant human M-CSF (R&D Systems, Abingdon, UK) at 37°C and 5% CO_2_. After two days, the cells were differentiated into mature OCs by further culturing them for seven days in medium containing 25 ng/ml of both M-CSF and recombinant human RANKL (R&D Systems, Abingdon, UK) and medium was changed twice.

### OB preparation

OBs were isolated from outgrowths from trabecular bone of patients undergoing hip replacement (ethical approval S-20110114 and informed consent) ^8^. In brief, trabecular bone was dissected into ∼5 mm pieces, which were cleaned twice by vigorous shaking in PBS. 5-6 pieces per well of a 12-well plate were cultured at 37 °C, 5% CO_2_ for around 3 weeks (i.e. roughly 80% confluency) in 2.5 ml BMOB media (Dulbecco’s Modified Eagle Medium (DMEM); Sigma-Aldrich, St Louis, MO) containing 2 mM L-glutamine, 50 µM ascorbic acid, 10 nM Dexamethasone, 10 mM glycerophosphate, and 10% FCS. Media were changed every 7 days. The cells were thereafter reseeded into T25 cell culture flask (2 x 10^5^ cells).

### Cell and bone surface labelling for time-lapse imaging in co-cultures

Bovine cortical bone slices (0.4 mm; BoneSlices.com, Jelling, Denmark) were labeled with rhodamine fluorescent dye (ThermoFisher) as described ^19^. OCs were detached from T25 flasks with accutase, centrifuged, resuspended in αMEM containing10% FCS, 25 ng/ml M-CSF, and 25 ng/ml RANKL, and reseeded in a 96-well plate containing labeled bone slices (100,000 cells per slice). F-actin in OCs was labeled ^19^ by incubating them for 5 h in 100 nM SiR-actin (excitation: 652 nm; emission: 674 nm) and 10 μM verapamil (both from Spirochrome, Stein am Rhein, Switzerland) at 37°C in 5% CO_2_. Two hours later, staining of OBs was initiated. OBs in T25 flasks were washed with PBS and incubated in 5 ml BMOB medium containing 5 µM Vybrant DiO (Invitrogen, Taastrup, Denmark). After 20 min, OBs were washed twice for 10 min with BMOB medium and once in PBS. After detachment with accutase, the OBs were centrifuged at 400 g for 5 min and the pellet was re-suspended to 2.5x10^5^ cells/ml in αMEM media containing 25 ng/ml M-CSF, no RANKL, no FCS. After 3 h incubation, the media of the OC-loaded bone slices were discarded and replaced by 200 µl fresh SiR-actin-Verapamil αMEM media and 50 µl labelled OB cell suspension (=12,500 cells), and incubated for 2 h. Inhibitors of CatK activity (ODN; 15 nM, 50 nM and T06; 300 nM, 500 nM) were added at this point in a 10 µl-volume. Efficacy of these inhibitors was tested previously (ODN: IC_50_ for collagen degradation 0.21 ± .05 μM , for OC resorptive activity 0.014 ± 0.004 μM, ^17^; T06: IC50 for collagen degradation 2.7 ± 0.2 μM ^30^, for OC resorptive activity 0.24 ± 0.06 μM^18^ (Supplementary Table S1). After these incubations, bone slices were transferred (inverted) to the Nunc Lab-Tek II chambered cover-glass (ThermoFisher Scientific) containing αMEM (25 ng/mL M-CSF, no RANKL, no FCS) with or without inhibitors. Time-lapse imaging was done using Olympus Fluoview FV10i microscope (Olympus Corporation, Shinjuku, Tokyo, Japan) at 5% CO_2_ and 37°C, using a 10x objective with a confocal aperture of 2.0 corresponding to a z-plane depth of 20.2 µm ^19^. The initial focus was set on the bone surface. Interval between frame recordings was ∼23.6 min and total recording was for 72 h (total 183 frames). At least 3 areas per bone slice were chosen for imaging and adjusted with Cy5 (EX645 nm, EM664nm), Rhodamine Phalloidin (EX558 nm, EM575nm) and DiO (EX491 nm, EM506nm) fluorescence settings. Neither SiR-actin, verapamil, nor DiO affected bone resorption.

### Analyses of time-lapse recordings for OC-OB co-culture

Olympus Fluoview 4.2 Viewer (Olympus Corporation) was used to optimize the intensities of different channels and exported into “.avi” format. The videos were analyzed with Fiji; ImageJ (NIH, USA) for different parameters ^16,19^. The duration of each OC or OB activity during or after cavity formation was measured by counting frames and converting counts into time. The number/frequency of simultaneous and consecutive OB visits to a given pit/trench occupied or not by an OC was also counted. Since SiRActin also stains somewhat the OBs, the circumference of the osteoblast is visible when using both the SiRActin (green) and the DIO (cyan) filter. OB interactions with resorbed cavities were defined by (i) more than 50% cell surface over/inside the excavation viewing via DiO(cyan) and rhodamine (red) filter and (ii) their occupancy over a period of more than one frame (23.6 min). Sometimes more than one OB was associated with an excavation. The number of frames showing association with OBs was determined and the average time of individual OBs residing in the excavation was calculated. We discriminated: (i) the duration of direct “OC-OB interactions” (i.e. during initiation and generation of the excavation), and (ii) the whole-time span of “OB association with the excavation” (i.e. from the start of rhodamine disappearance from the bone surface until the end of the culture) which thus includes both the duration of OC-OB interaction and the time span after the passage of the OC. The presence of OBs in OC-positive and OC-negative pits/trenches was evaluated at three time points (24 h, 48 h, and 72 h) and the average was used to determine the percentage of pits or trenches associated with OBs. OCs initiate resorption by forming a circular F-actin ring (pit), which changes into crescent shape when pits extend into trenches. The appearance of an actin ring was set up to define OC-OB association when resorption starts, and the transition from ring to crescent shape to discriminate between OB association to pits and trenches.

### Quantification of OB-interactions with excavations at the end of cultures

These interactions were evaluated both in OC-OB co-cultures and in cultures of OBs seeded on pre-resorbed bone (i) For co-culture experiments, mature OCs were seeded at a density of 100,000 cells per bone slice in a 96-well plate and incubated at 5% CO_2_ and 37°C in αMEM (25 ng/ml M-CSF, no RANKL, no FCS). After 2 h, 12,500 OBs per bone slice were added. After 72 h co-culture in the presence or absence of CatK inhibitors, ALP-activity was measured in conditioned media by using a colorimetric assay (Sigma-Aldrich, St Louis, MO) and normalized to live cell numbers. C-telopeptide of type I collagen (CTX-I) concentration in culture media was measured by using an ELISA kit (MyBioSource, San Diego, CA, US). Bone slices were washed in PBS, fixed in 4% formaldehyde. OCs were visualized through TRAcP activity using leukocyte (TRAcP) kit (Sigma-Aldrich, St Louis, MO) and OBs were subsequently visualized using leukocyte alkaline phosphatase kit (Sigma-Aldrich, St Louis, MO) (ii) For culturing OBs on pre-resorbed bone slices, OCs were first cultured on bone slices for 72 h in the same conditions as for the co-cultures. Thereafter, bone slices were incubated in H_2_O and stripped using a cotton stick to lyse the OCs. Subsequently, 12,500 OBs were seeded on each pre-resorbed bone slice and cultured for 72 h in αMEM (25 ng/ml M-CSF, no RANKL, no FCS), in a 96-well plate at 5% CO_2_ and 37°C. Thereafter, OBs were visualized using the Leukocyte Alkaline Phosphatase Kit while excavations were visualized using toluidine blue staining. Possible remnants of OCs in excavations were assessed by TRAcP staining. Cell survival was assessed in parallel 72-h co-cultures or consecutive mono-cultures, by assessment of cell numbers and metabolic activity using CellTiter-Blue Viability Assay (Promega, Madison, USA).

### Western blot analysis of cell lysates from OC-OB co-cultures

After the culture, cells adhering on the bone were scraped in lysis buffer (RIPA, Millipore, Sigma) containing a protease inhibitor cocktail (Sigma-Aldrich, St Louis, MO). Proteins were quantified using (Bio-Rad kit (Richmond, CA, USA). 10 μg of protein per condition were subjected to SDS-PAGE (15%) and transferred onto nitrocellulose membrane using wet blot method (Bio-Rad) at 100V for 2 h at 4°C. After blocking in 3% (weight/volume) skimmed milk for 3 h, the membrane was incubated in TBST (Tris-buffered saline with 0.05% Tween-20) overnight at 4°C for detection of RANK (MAB6263-SP; R&D systems, Minneapolis, USA), ALP (AF2909-SP; R&D systems), COL1 (ab34710; Abcam, Cambridge, UK), MMP 13 (ab39012; Abcam), CatK (MS2),) and β-actin (ab8227, Abcam) (primary antibody dilution; 1:1,000). Membranes were washed 3 times with TBST buffer for 15 min and incubated in TBST buffer at room temperature for 1 h in HRP-conjugated secondary anti-IgG antibody (W4018, Promega, Madison, USA), at 1:2,500 dilution. Membranes were washed with TBST. Proteins were visualized with enhanced chemiluminescence assay kit (GE Healthcare Life Sciences, Piscataway, NJ, USA) using GE ImageQuant LAS 500 Imager (GE Healthcare Life Sciences). Protein levels were evaluated through densitometry (ImageJ software) and normalized to actin.

### Microchemical analysis of resorption cavities

Four types of experiments were performed :(i) mono-cultures of OCs^17^, (ii) OC-OB co-cultures, (iii) cultures of OBs on bone slices previously subjected to OC cultures, and (iv) cultures of OBs on bone slices previously subjected to OC-OB co-cultures. These cultures were run as aforementioned for 72 h in untreated, T06, and ODN-treated conditions. Thereafter, bone slices were incubated in dH_2_O for 10 min, wiped smoothly with cotton stick to remove the cells, fixed in 2.5% glutaraldehyde and prepared for scanning electron microscope (SEM) and energy dispersive X-ray spectroscopy (EDAX) at voltage of 8 kV and beam current of 13 nA ^17^. Spectrum within 5µm^2^ region of interest (ROI) (5-10 regions per pit in each condition for 25 pits or trenches) were analyzed in three experiments including cells from different donors. The atomic percentage of elements present in excavations, was determined by EDAX, TEAM Pegasus analysis system. Secondary electron (SE) imaging was used to observe structural variations within excavations and backscattered-electron (BSE) imaging for topography, crystalline structure, and microchemical analysis.

### Statistical analysis

Time-lapse studies included four independent experiments from different human donors in each condition (40 time-lapse videos). Similarly, four independent 72 h endpoint experiments from different donors in CatK inhibited conditions were conducted to observe OBs activity, differentiation, and their preference towards excavation-type. For EDAX analysis 3 independent experiments per condition from different donors were conducted. All statistics shown in graphs were performed using GraphPad Prism software, version 9 (GraphPad software, San Diego, CA, USA), and SigmaPlot software (SPSS Inc.). P < 0.05 was defined as the level of significance. Data sets were analyzed for normal distribution following a Gaussian distribution; D’Agostino & Pearson omnibus test, and statistical tests were adapted accordingly. Data are presented as median. Kruskal–Wallis test, two tailed; Dunn’s multiple comparisons test was used for most of the analyses. More details on the employed statistics are given in the figure legends.

## RESULTS

### Association of OBs with pit and trench resorption events

First, we performed time-lapse analysis of OC-OB co-cultures on bone slices, and observed for 72 h whether OBs would respond to bone resorption events and whether this response would differ depending on the resorption mode (Figure 1A-C, video 1). As reported^16,19^, OCs start resorbing in pit-mode, and most of them thereafter shift to trench-mode leading to resorption for longer periods and over longer distances. We observed frequent OB-OC-interactions at the initiation of resorption, as soon as the actin ring, typical of pit-mode, appears and these interactions continue until the transition from pit- to trench-mode. However, OB-OC interactions are lost when the OCs shift into trench-mode as well as during trench extension. Both average occupancy and individual time of OBs in pits are higher than in trenches (Figure 1D, E, Video 1). The frequency of OB visits to pits is higher compared to trenches, irrespective of whether an OC is present or not (Figure 1F). Timelines for OB-interactions with pits or trenches (Figure 1G) show that OBs interact with trenches primarily at the beginning of resorption. In contrast, OBs interact with pits at different intervals of the observation period. The OB occupancy time in pits is higher than that of OCs, whereas the opposite is true for trenches (Figure 1G).

**Figure 1.**
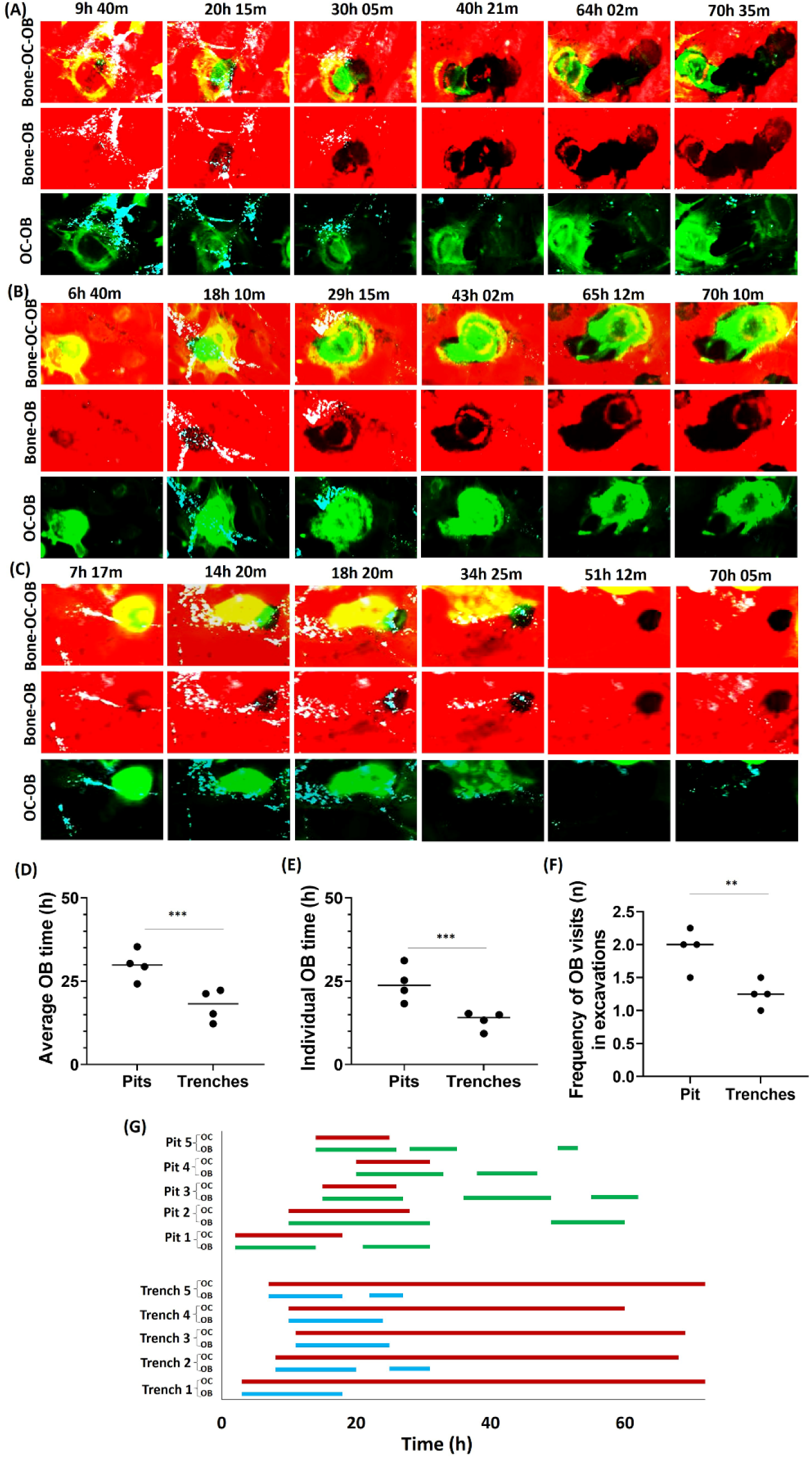
Effect of the pit/trench resorption mode on the association of OBs with resorption events. (A-C) Selected time-lapse images of OCs co-cultured with OB forming trenches (A, B, video1) and pits (C, video 1) during 72 h of co-culture. OCs were stained for actin by using phalloidin (green), OBs were stained with vybrant DiO cell-labeling solution (cyan), and bone surface was stained with rhodamine (red). Resorption of bone surface by OCs is shown by black trench/pits imprints. (D) Average OB time (+/-OCs), and (E) individual OB time (+/-OCs) associated with particular type of excavation formation confirms OBs preference to the pits. (F) Frequency of OB visits (total number of OBs resided simultaneously/consecutively) to pits and trenches during this observation confirms more visits of OBs to pits. Data shown are (n: 4 donors; with 8 repeats). The median proportions obtained in each experiment are shown as bars. We analyzed between 40-120 cells activities per condition per experiment for each of the donors). Statistics: Mann-Whitney test (ns: not significant; **P < 0.01; ***P < 0.001). (G) Graphical distribution of OB-OC-excavation-interactions over 72 h revealed that OBs reside in the beginning of resorption event; however, pits were often occupied consecutively by OBs throughout monitoring time compared to trenches which lacks OB interaction during elongation (track/pattern of 5 OB/OC residing in trenches and pits from beginning till end of co-culture). OCs distribution shown in red, OB distribution in pit shown in green, and OB distribution in trenches shown in blue.

### CatK-inhibition enhances the association of OBs with pit and trench resorption events

OC-OB interactions at the onset of bone resorption and their loss during trench expansion under control conditions (Figure 1, Video 1) are confirmed in Figure S1A and Figure 2. At low concentrations of T06 (300 nM) and ODN (15 nM), OCs still generate small trenches and more pits. Interestingly, compared to control conditions, these trench resorption events show longer and more frequent interactions with OBs (Figure S1B-C, Figure 2, Video 2). Maximum CatK inhibition with T06 (1µM) and ODN (50 nM) prevents the switch from pit to trench mode, thereby inducing mostly pit formation. CatK-inhibited OCs show longer involvement in pit formation, which corroborates with longer OB interactions with OCs and the generated pits (Figure S1D-F, Figure 2 I, Video 3).

**Figure 2.**
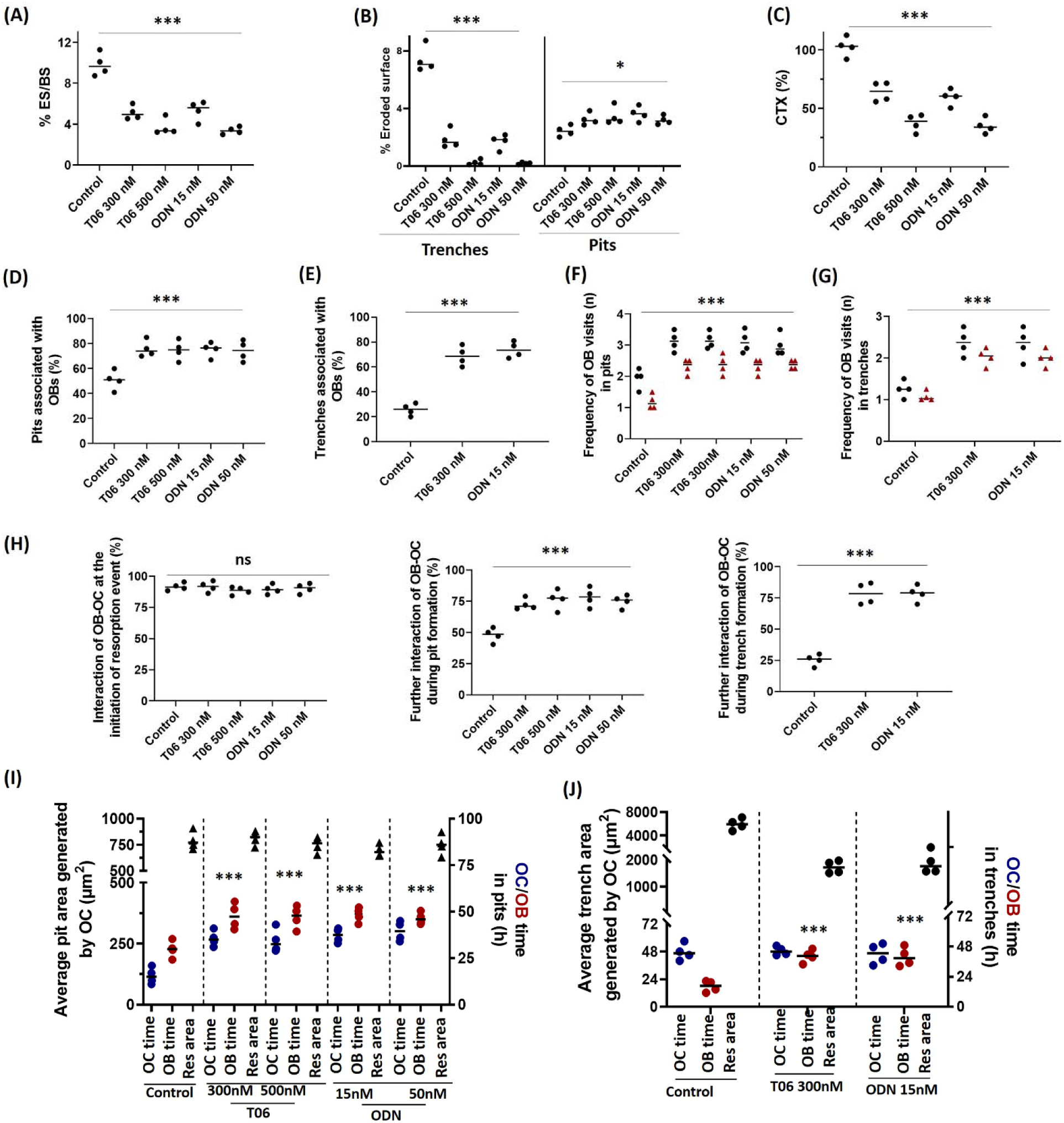
Quantitative analysis of CatK inhibitor effects on the association of OBs with resorption events. (A) Effect of T06 and ODN on the % eroded surface, (B) % of eroded surface in terms of trenches and pits, and (C) CTX release in the conditioned media after 72 h. (D-E) Percentage of pits or trenches (+/-OCs) associated with OBs and their preference for pits over trenches in the presence and absence of CatK inhibitors. (F-G) The frequency of OBs visits to pits and trenches during 72 h-co-cultures in +/- of CatK inhibitors. Data shown in black is for total number of OB visits and data shown in red is the number of simultaneous occupancies by OBs. (H) The percentage (%) of OB-OC interaction was highest when OC initiates resorption and decreased at later stages for both pits and trenches generation. However, at these latter stages, OC-OB interactions remain frequently seen if CatK inhibitors are present. (Statistics: Kruskal–Wallis test, two tailed (ns: not significant; *P < 0.05; ***P < 0.001); Dunn’s multiple comparisons test (ns: not significant; *P < 0.05; ***P < 0.001) compared with untreated control. (I) The average resorbed area generated by individual OCs in form of pits and OBs interaction with these pits along with OC/OB interactions are shown in control and CatK inhibited conditions. OC/OB and OBs interaction with these pits was longer with CatK inhibition. (J) Time spans of OCs to generate individual trenches (average resorbed area generated by individual OCs in form of trenches) and the duration of trenches associated with OBs are compared in +/- of CatK inhibitors. Sample size: Control: n=4 donors; 300 nM T06: n=4 donors; 500 nM T06: n=4 donors; 15 nM ODN: n=4 donors; 50 nM ODN: n=4 (2 donors). For each donor, 2-3 replicate experiments on individual bone slices were analyzed for all conditions. The median obtained in each experiment are shown as bars. We analyzed between 40-120 OC-OB activity per condition per experimental repeat for each of the donors). Statistics: Kruskal–Wallis test, two tailed (ns: not significant; ***P < 0.001); Dunn’s multiple comparisons test (ns: not significant; ***P < 0.001) compared OBs occupancy time with untreated control.

Figure 2 shows a quantitative analysis of the features observed in Figs S1. Inhibition of CatK by both T06 and ODN strongly reduces bone resorption, as indicated both by the eroded surface and CTX release (Figure 2A, C). Of note, in untreated conditions, OCs generate ∼28% pit-eroded surface and ∼72% trench-eroded surface (Figure 2B). Maximum CatK-inhibition completely abolishes trench-eroded surface, but increases pit surface by 20% compared to control (Figure 2B). The OB-occupancy of pits is significantly higher than that of trenches and OB-occupancies both of pits and trenches are increased in response to CatK inhibition (Figure 2D, E; Figure S2A, B). Under untreated conditions, the frequency of OB visits to pits is significantly higher than that to trenches. These frequencies (including simultaneous visits) increase with CatK inhibition (Figure 2F, G). At the initiation of resorption, when the actin ring appears, almost all resorption events show OC-OB interactions, whether CatK is inhibited or not. However, as soon as resorption proceeds and rhodamine removal is visible, OC-OB interactions drop to about 50% in the pit resorption mode and to about 25% when switching to the trench resorption mode. Still, both these percentages remain significantly higher when CatK is inhibited (Figure 2H). Furthermore, inhibition of CatK significantly increases the overall average OB exposure time to pits and trenches (including duration of OC-OB interaction and time span after the passage of the OC (Figure 2I, J). Under untreated condition, OCs are engaged for longer time in trench formation compared to pit formation. The effect of CatK inhibition on the duration of resorption events in co-cultures is just as reported for monocultures^16^: pit-forming OCs remain associated with the resorption cavity for longer times (Figure 2I), but trench-forming OCs continue elongating the trench at a slower rate, thus resulting in shorter trenches (Figure 2J). Frequency and duration of OB occupancy in pits was higher despite trenches represent a larger resorbed area (Figure 2I). Interestingly, the duration of OB association with pits is always longer than that of OCs, whether CatK inhibitors are present or not (Figure 2J). In contrast, the duration of OB association with trenches is shorter than that of OCs in control conditions, but equals that of OCs subjected to CatK inhibitors (Figure 2J).

### OBs remain associated to “stagnating” OCs for as long as the OC is associated with the resorption cavity

We previously reported that ∼25% of the resorbing OCs in monoculture show a stagnating activity when treated with either T06 (500 nM) or ODN (15 nM or 50 nM)^16^. Stagnating activity is characterized by persistent engagement of the OC at the same excavation, moving erratically while orienting the sealing zone and ruffled border in multiple successive directions, and decelerating resorption. CatK inhibitors also induce this typical stagnating activity in OCs co-cultured with OBs (Figure S3A-C; video 4). Therefore, we analyzed whether stagnating activity affects the association of excavations with OBs and OCs. CatK inhibition prolongs the time an OC remains associated with a pit (also shown in Figure 2), but this association time is further prolonged in case of stagnating activity (Figure S3D). Interestingly, OBs remain associated to the resorption event site as long time as the stagnating OC persists at the same excavation (Figure S3 D).

### CatK inhibition enhances the maturation of OBs in resorption cavities

The above data show that the way bone is resorbed (i.e. pit vs. trench resorption mode and low vs. strong collagenolysis/CatK activity) matters for OB recruitment into the resorption cavities. Next, we investigated whether the way of resorption changes OB maturation. First, we found that the proportion of ALP^+^ OBs associated with pits at the termination of co-cultures is higher compared to that associated with trenches (whether or not OCs sit on the cavity) (Figure 3 A,B). Furthermore, CatK inhibition enhances the association of ALP^+^ OBs to both pits and trenches (Figure 3B). A quantification of OC and ALP^+^ OB co-localization confirmed more OC-OB interactions in resorption cavities after CatK inhibition (Figure 3C). These findings are thus just as shown for the Vybrant-labeled OBs in the time-laps experiments (Figure 2). Note also that T06 and ODN do not affect cell viability whether in co- or mono-culture (Figure S4). Interestingly, both CatK inhibitors significantly enhance ALP-activity in the conditioned media (Figure 3D). Furthermore, Western blot analyses of cell lysates from OB-OC co-cultures, showed that T06 and ODN increase ALP, COL1, MMP13, and CatK protein levels but reduce RANKL protein levels after 72 h co-culture. (Figure 3E-F).

**Figure 3.**
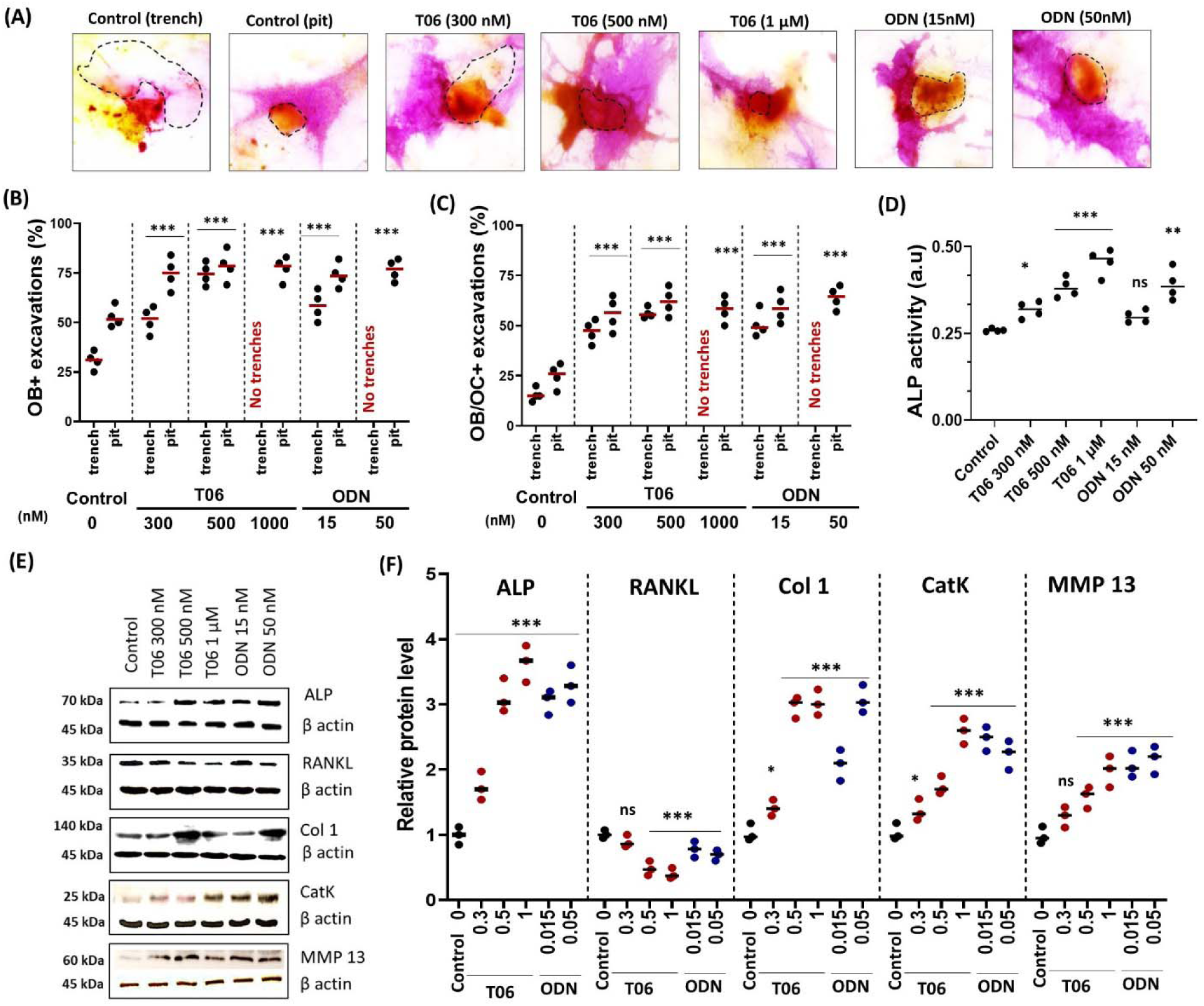
Prevalence of ALP^+^ OBs in resorption cavities and level of osteogenic markers after a 72 h-coculture of OBs and OCs in the presence and absence of CatK inhibitors. (A) Representative images of TRAP-stained OCs (orange) and ALP-stained OBs (purple) under control, T06 and ODN treated conditions. The segmented outline (black dotted) shows the resorption excavation generated by OCs. (B) The quantification of ALP^+^ resorption cavities (+/- OCs) during 72 h of OC-OB co-culture indicated the preference of OBs towards pits compared to trenches when normalized with total number of pits and trenches. (C) The quantification of ALP^+^ resorption cavities with OCs (+ OCs). Statistics: Kruskal–Wallis’s test, two tailed (*P < 0.05; ***P < 0.001); Dunn’s multiple comparisons test (*P < 0.057; ***P < 0.001) compared to the untreated control. (D) ALP activity quantification in the culture supernatant showed significantly enhanced activity under CatK inhibition (Kruskal–Wallis’s test, two tailed (ns: not significant; *P < 0.05; **P < 0.01; ***P < 0.001). Sample size (n=4 donors) for control, T06 (300 nM, 500 nM, 1 µM), and ODN (15 nM, 50 nM). For each donor, 3 replicate experiments on individual bone slices were analyzed for all conditions. The median obtained in each experiment are shown as bars. We analyzed between 450-800 OC-OB activity per condition per experiment for each of the donors). (E) Western blot analysis of ALP, RANKL, collagen (COL1), CatK and MMP 13 in control, T06 (0.3, 0.5, and 1 µM), ODN (15 and 50 nM) conditions and (F) their quantification (n=3 donors). Statistics: Kruskal–Wallis’s test, two tailed (ns: not significant; **P < 0.01; ***P < 0.001); Dunn’s multiple comparisons test (**P < 0.02; ***P < 0.001) compared to untreated control.

Seeding OBs (in the absence of OCs) on pre-resorbed bone allowed to evaluate whether the type of eroded surface independently of the presence of OCs - would influence the degree of association of OBs with different types of eroded surfaces. Figures 4 A-D show images of OBs in contact with pits or trenches. The relative prevalence of OB-positive resorption cavities showed the OBs select pits rather than trenches (Figure 4E), as they also did in OC-OB co-cultures (Figure 3B). Thus, the eroded matrix of pits is more attractive than that of trenches. Furthermore, CatK inhibition during the OC preculture renders pits and trenches even more attractive (Figure 3B), but addition of CatK inhibitors during the OB culture did not have any effect indicating that both inhibitors do not interfere with OB activity (data not shown). Of note, the higher the prevalence of OB-positive excavations, the higher the ALP staining of the OBs associated with these excavations (Figure 4E-F).

**Figure 4.**
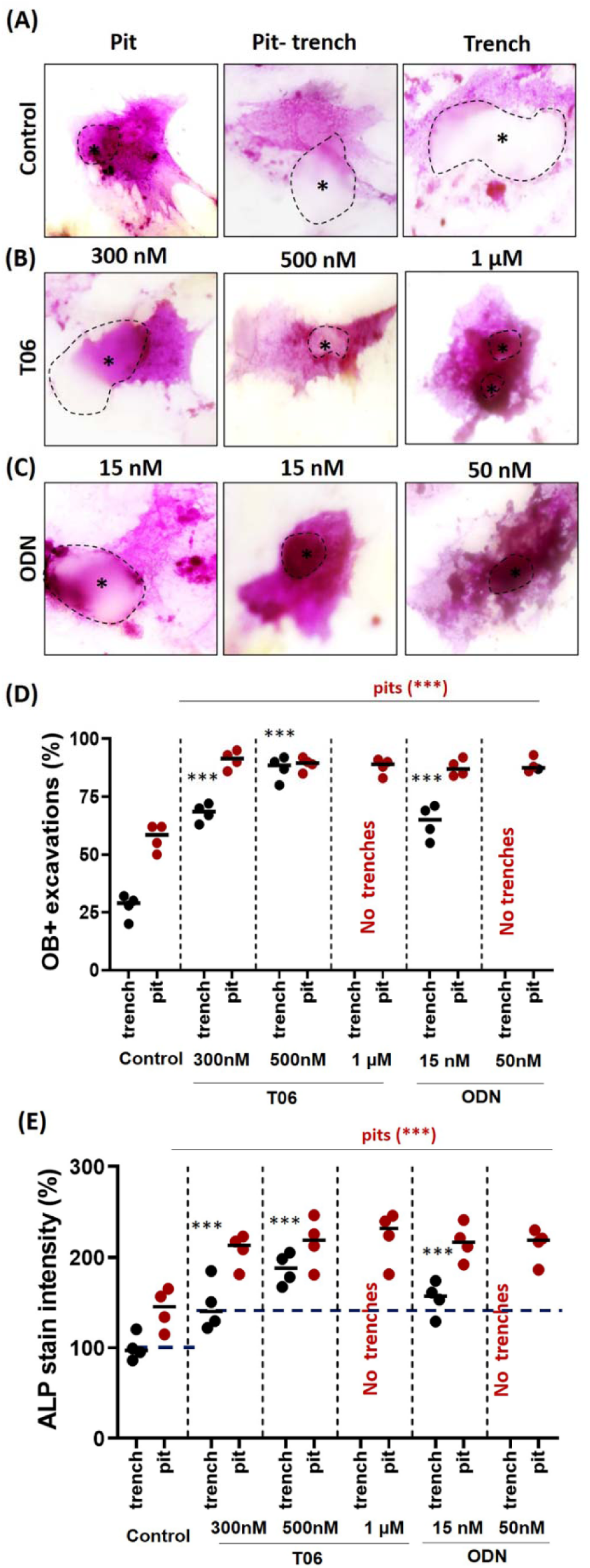
Comparative response of OBs to pits and trenches generated during a preculture with OCs subjected or not to CatK inhibition. OB cultures on bone slices which have been pre-treated with OCs subjected or not to CatK inhibitors: prevalence and ALP activity levels in OBs associated with pits and trenches: Bone slices were submitted to OC resorption for 72 h according to the standard protocol under control, T06 (0.3, 0.5, and 1 µM), and ODN (15 and 50 nM), conditions. After stripping of OCs, OBs were seeded on these pre-resorbed bone slices and cultured for another 72 h (CatK inhibitors -/-). (A-D) Representative images of ALP-stained OBs (purple) after this culture on the bone slices pretreated as indicated. Asterisks and dotted outline highlight the position of resorption cavity. (E) Prevalence of ALP^+^ excavations according to whether they were generated as pits, trenches, and in the presence or absence of CatK inhibitors. Pits and trenches generated in the presence of CatK inhibitors are more likely to attract OBs. (F) Levels of ALP activity in OBs according to whether they are associated with pits and trenches that had been generated in the presence or absence of CatK inhibitors. Horizontal dotted lines represent the average staining intensity of OBs on the non-resorbed bone surface under different conditions. They showed less ALP-activity compared to resorbed excavations. Sample size (n=4 donors) for control, T06 (300 nM, 500 nM, 1 µM), and ODN (15 nM, 50 nM). For each donor, 3 replicate experiments on individual bone slices were analyzed for all conditions. The median obtained in each experiment are shown as bars. We analyzed between 500-900 OB activities per condition per experiment for each of the donors). Statistics: Kruskal–Wallis’s test, two tailed (**P < 0.01; ***P < 0.001); Dunn’s multiple comparisons test (**P < 0.01; ***P < 0.001) compared to the untreated control.

### Ultrastructural and microchemical analysis of the excavations in OC-OB co-culture

Next, we analyzed the physico-chemical characteristics of excavations by SEM and EDAX for parameters reported to favor bone formation at reversal surfaces, such as smoothness and mineralized surfaces devoid of collagen remnants ^13,32^. SEM shows less collagen and more mineral clusters in pits generated by OCs co-cultured with OBs, compared to pits generated by OCs alone (Figure S5 A-B). Similar effects are observed when OBs are cultured on pre-resorbed bone (Figure S5C). An even higher increase in mineral deposition is noted when OBs are cultured on pre-resorbed bone during OC-OB co-cultures (Figure S5D). EDAX analysis of the bottom of pits confirms increased calcium and phosphorous (reflecting mineral) and decreased carbon and nitrogen (reflecting primarily collagen) in pits subjected to OBs (Figure S5E). Changes in oxygen are less informative as both collagen and mineral contain oxygen. Next, we evaluated whether CatK inhibitors change the mineralization characteristics of the eroded surfaces of pits and trenches generated in OC mono-cultures or OC-OB co-cultures. Figures 5A, B confirm the more complex topography and mineral clusters in pits generated in the presence of OBs. Clusters are even more abundant under CatK inhibition. Spectrometry measurements are in accordance with these morphological assessments (Figure 5C): pit surfaces in monocultures show a high carbon peak and low calcium and phosphorous peaks, whereas the carbon peak is lower and the calcium and phosphorous peaks are higher for pit surfaces in co-cultures. These respective changes are even larger in the presence of CatK inhibitors (Figure 5C, Figure S6A). Unlike pits and as already mentioned, trenches generated in control OC monocultures are devoid of undegraded collagen, but do show undegraded collagen under CatK inhibition. Less of this undegraded collagen and mineral clusters is seen in co-cultures (Figure 5D). Again, these visual observations are confirmed by the spectrometric assessments shown in Figure 5E and Figure S6B.

**Figure 5.**
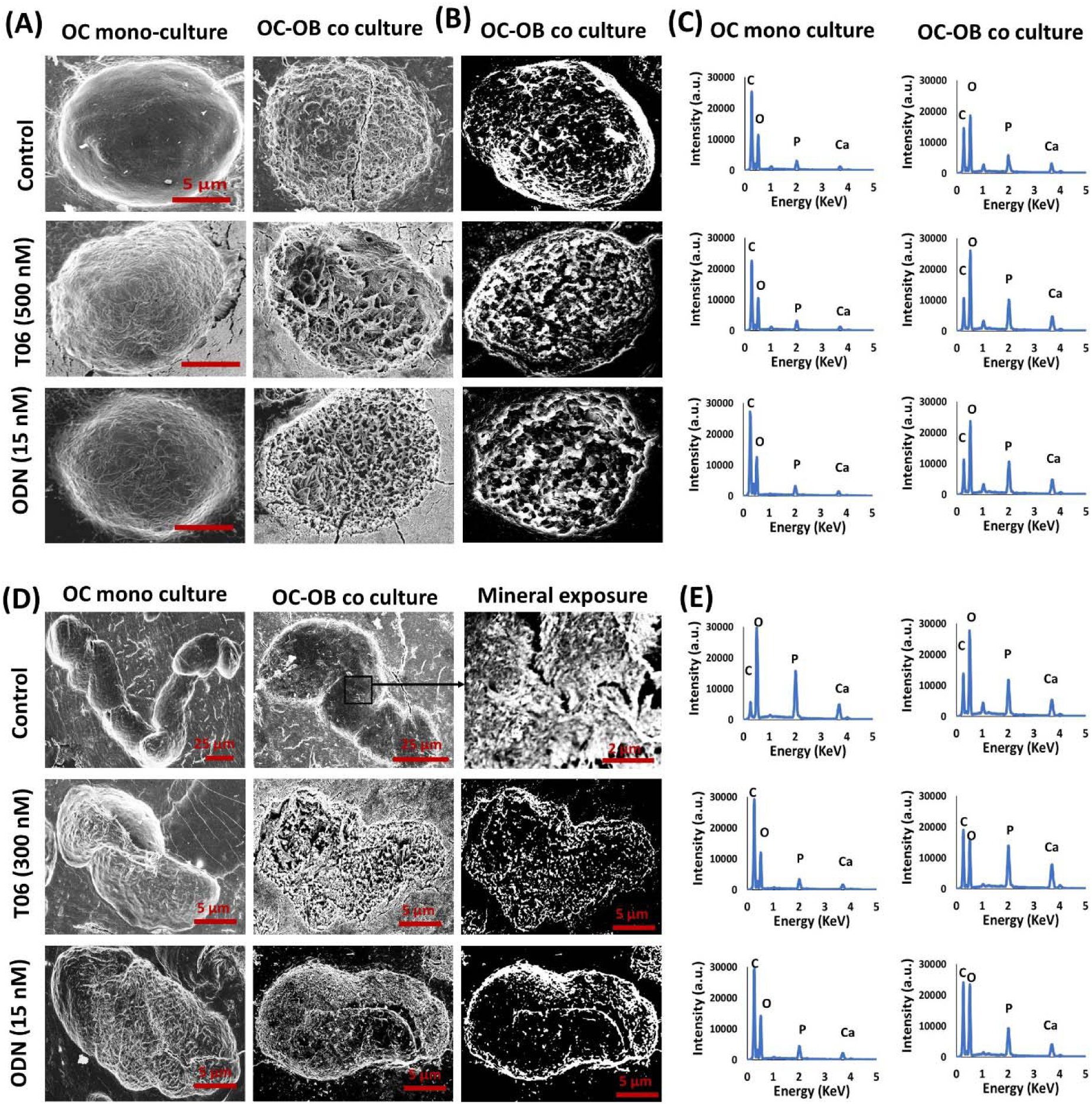
Mineral exposure in resorption cavities during OC-OB co-cultures. (A) SEM micrographs obtained through secondary electron imaging show pits generated by OC mono-cultures and in OC-OB co-culture under the indicated culture conditions. The topography of the pit surface generated during co-culture is more complex than that generated in mono-culture. (B) Micrographs obtained through backscattered electron imaging clearly confirm this difference. Scale bar: 5 µm for pits in each condition. (C) Energy dispersive X-ray spectroscopy spectra show a prevalence of carbon in pits of mono-cultures which reflects non-degraded collagen, whereas a higher prevalence of P and Ca is seen in pits generated in OC-OB co-cultures compared to those generated in OC mono-cultures, and even more so in co-cultures performed in the presence of CatK inhibitors. (D-E) Physical analysis of trenches. (D) Micrographs obtained through backscattered electron imaging show that trenches generated in mono-cultures under control conditions do not show undegraded collagen, but trenches generated under CatK inhibition do so. (E) Energy dispersive X-ray spectroscopy spectra clarify these observations. They show a low C peak and high Ca and P peaks in trenches generated in OC mono-cultures under control conditions (reflecting a mineralized surface without non-degraded collagen), whereas the reverse is true in trenches generated in OC mono-cultures performed under CatK inhibition (thus reflecting non-degraded collagen on the resorbed surface). However, if CatK inhibition was performed in co-cultures, Ca and P peaks became higher, thus reflecting exposure of mineral at the ES. Minor traces of impurities were observed and normalized in the quantification. C: carbon; O: oxygen; P: phosphorous; Ca: calcium.

## DISCUSSION

CatK inhibitors reduce bone resorption mainly by prompting OCs to make shallow resorption pits rather than long and deep resorption trenches ^14,16–18,20,22,30^. Here, we show a greater association of OBs with pits compared to trenches, thereby demonstrating that low CatK activity levels favor the occupancy of eroded surfaces by OBs (Figure 6). Our evidence is based on (i) the greater number and duration of OB visits to pits compared to trenches in OC-OB co-cultures analyzed in real time, (ii) the higher proportion of OB-positive pits compared to OB-positive trenches at termination of a 3-day co-culture, as well as (iii) the greater pit occupancy by OBs seeded on bone pre-resorbed by OCs. This OC-OB proximity allows OBs to contribute to resorption by degrading demineralized collagen as evaluated by SEM and spectrometry, and to differentiate faster as shown by differentiation markers. These diverse data all indicate that pits and trenches may differently influence the preparative steps of bone formation during bone remodeling.

**Figure 6.**
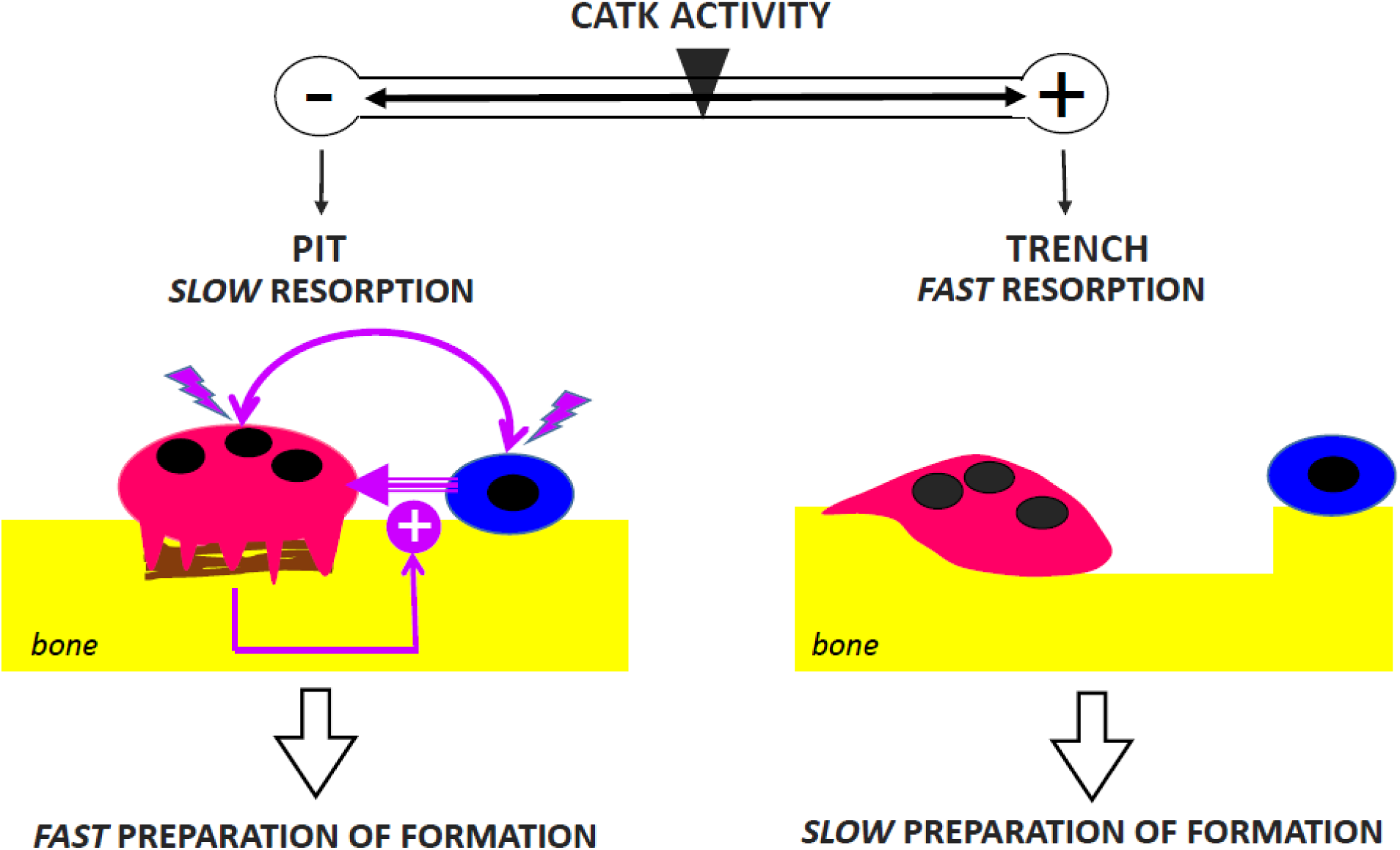
Low CatK activity promotes OC-OB proximity and fast preparation of bone formation. Low CatK activity leads to accumulation of demineralized collagen (brown) beneath the ruffled border of the OC (red), so that the OC soon becomes inactivated and can only generate shallow pits. According to our model, demineralized collagen attracts osteoblast lineage cells (blue), thereby favoring the OC-OB interactions that are required to induce bone reconstruction (see discussion). In contrast, high CatK activity ensures complete removal of collagen by the OC, thereby allowing continuous resorption in the form of long and deep trenches. Trenches are devoid of collagen and therefore less attractive for OBs (see discussion).

### Low levels of OC collagenolysis enhance the prevalence of OBs on eroded surfaces

The intriguing question is, which mechanism relates the prevalence of OBs at resorption events to CatK activity levels and to pit vs. trench resorption mode. First of all, it is clear that CatK inhibitors do not affect the association of OBs with OCs before resorption has started: at the time of activation, almost all OCs are associated with OBs whether CatK inhibitors are present or not (Figure 2H). This association may be explained by the absence of addition of RANKL to the culture fluids: activation of an OC may only occur if the OC happens to be next to a RANKL- expressing OB. This interpretation is supported by the strong inhibition of resorption by OPG in this OC-OB co-culture model^8^.

It is only after bone resorption has started that the prevalence of OBs becomes sensitive to CatK inhibitors. In OC monocultures, almost all resorption events start in the pit mode ^16,19^, and this holds true in the presence of OBs (present study). Pit formation consists of the invasion of the OC ruffled border perpendicularly into the bone matrix, and demands a strict synchronization of the speed of demineralization and degradation of the demineralized collagen ^15^. Thus, low CatK activity leads to the accumulation of demineralized collagen between the ruffled border and the bone matrix - which prevents further penetration of the ruffled border into the bone. So, OCs make round pits containing demineralized collagen (Figure 6). In contrast, high CatK activity enables full collagen degradation and thereby a deep cavitation, wherein the sealing zone can move to re-orient the ruffled border against the cavity wall and switch the OC into the trench resorption mode (Figure 6). Pits are thus clearly different from trenches regarding the presence of demineralized collagen ^17,20,33^. Since collagen fibers exert very strong haptotaxis on OBs compared to other bone matrix proteins in *in vitro* assays ^34^, we believe that the presence of collagen in pits, but not in trenches, explains the larger OB occupancy of pits compared to trenches. This role of collagen in OB recruitment is further supported by more and longer OB visits to pits generated in the presence of CatK inhibitors (Figure 2). It is also corroborated by the fact that the few trenches that could be generated, despite the presence of CatK inhibitors, show collagen fibers (Figure 5) and in these situations (but not in control conditions) OBs remain associated to the trench as long as the OC (Figure 2J, 1G). Even the frequency of simultaneous OB visits is increased (Figure 2F, G), which is of interest because OB density on eroded surfaces is critical for initiation of osteogenesis ^35^.

Of note, also *in vivo* observations support a role of collagen in homing OBs onto eroded surfaces: (i) EM pictures of human bone show OB lineage cells (reversal cells) interacting with collagen at the OC-bone interface ^13,34^. (ii) Mouse bone cultures in the presence of CatK inhibitors show abundant OB lineage cells enwrapping demineralized collagen fibers in resorption cavities ^13^. (iii) Bone sections of pycnodysostotic patients who lack CatK activity due to a mutation, show reversal cells enwrapping demineralized collagen left over in the resorption lacunae^13^. (iv) In ovariectomized rabbits, CatK inhibition was associated with both accumulation of demineralized collagen in the sub-OC resorption zone and a higher density of OB lineage cells on eroded surfaces ^36^.

One may of course think of alternative mechanisms explaining how CatK inhibitors affect the presence of OBs. For example, one could speculate that CatK inhibitors inhibit the degradation of factors such as TRAcP, IGF, BMP2, or TGF-β^17,18,37^ likely to affect homing of OBs on eroded surfaces ^1^. However, this speculation would only have value for ODN, which blocks the active site of CatK and thus CatK activity on all its substrates - but not for T06, which is an ectosteric inhibitor that specifically only blocks collagen degradation and leads here to same effects on the OBs as ODN ^18,30^. One may also think of the release of osteoclastic soluble agents, such as S1P, PDGF-BB, or active IGF, which are known to be upregulated by CatK inhibitors and to promote chemotaxis and proliferation of OB lineage cells ^37–39^ . However, pit-eroded surface is shown more attractive for OBs than trench-eroded surface irrespective of the presence of an OC: (i) the duration of OBs in pits is always longer than that of OCs (Figures 1, 2). (ii) OBs seeded on bone pretreated by OCs preferentially home in pits compared to trenches (Figure 4). (iii) Histology pictures of bones cultured in the presence of CatK inhibitors show OB lineage cells enwrapping collagen in resorption lacunae vacated by the OC ^13^. Thus, we believe that distinct characteristics of the eroded surface itself - and importantly demineralized collagen - significantly influence the presence of OBs on eroded surfaces. The importance of the OC signals “saved” on the eroded surface is in line with the need of OCs to communicate with OBs after their departure ^1,32^.

### Low levels of OC collagenolysis may be compensated by collagenolysis by OBs adjacent to OCs

An important observation is that OBs homing on the eroded surface seem to degrade collagen: pits are mostly filled with demineralized collagen after an OC mono-culture, but subsequently culturing OBs on these bone slices leads to removal of this collagen (Figure S5), thereby confirming earlier data ^23,40,41^. Likewise, pits generated in OC-OB co-cultures expose less collagen and more mineral than pits generated in OC monocultures (Figures 5, 6; Figure S5). Furthermore, the presence of CatK inhibitors in the co-cultures renders more thorough clearance of collagen by OBs. We show that this may be due to induction of the collagenase MMP13 in the culture settings containing either T06 or ODN (Figure 3E, F). MMP13 has also been shown to be upregulated by CatK inhibitors in *in vivo* studies ^42^ and is expressed by OBs next to OCs in our co-culture model ^8^. This OB-mediated collagen degradation contributes to OC bone resorption, since bone resorption is decreased by an MMP inhibitor in the present co-culture model, but not in mono-cultures of OCs ^8^.

Of note, observations on bone tissue support this view: (i) MMP13-positive OBs next to OCs were repeatedly reported (i.e. reversal cells on the bone matrix and canopy cells along the bone marrow) ^2,3,43,44^; (ii) MMP13-immunoreactivity was even detected between the ruffled border and the bone matrix in sections of mice and rabbit bone ^43,45^; (iii) an MMP inhibitor induces an accumulation of demineralized collagen between the ruffled border and the bone matrix while also inducing a corresponding decrease of bone resorption ^46^. Importantly, we show here that this OB contribution to collagenolysis is not enough to fully compensate a complete absence of CatK activity, as full CatK inhibition leads to stagnating resorption despite of the continuous association of OBs to OCs (Figure S3).

### Low levels of OC collagenolysis promote OB differentiation

Eroded surfaces generated under low CatK activity also promote OB maturation as shown by the relative levels of ALP associated to pits and trenches generated in co-cultures (Figure 3) or OC precultures (Figure 4). Furthermore, ALP increases a 3 to 4-fold in the conditioned medium and in the cell layer, and collagen increases a 3 to 4-fold in the cell layer in response to CatK inhibitors (Figure 3). In contrast, RANKL, which is known to decrease upon OB differentiation ^4,6^, is down-regulated in the presence of CatK inhibitors (Figure 3 E,F). It is tempting to relate this faster maturation to the removal of collagen remnants by MMP13, since this removal is mandatory for new matrix deposition on eroded surfaces ^13,41^. MMP13 appears thereby a pivotal player in bone remodeling: both completing resorption and facilitating OB differentiation.

### General *in vivo* relevance of the present OC-OB co-cultures

The above overview shows that our model of “OC-OB co-culture on bone slices” reproduces major OC-OB interactions of the bone remodeling cycle, starting as a pro-resorptive partnership and shifting to an osteogenic partnership ^1^. The importance of collagen and its degradation for linking OC and OB activities was already brought to mind by studies performed in the complexity of bone tissue ^13,34,36,38,39^. The present simplified model allows to demonstrate the intrinsic ability of collagen and its degradation by CatK and MMP13 to promote the interaction of OCs and OBs and thereby prepare for osteogenesis on eroded surfaces.

*In vivo*, OB lineage cells form continuous cell sheets at the bone surface (bone lining cells, reversal cells, bone marrow envelope, and canopy cells), thus providing bone formation potential very close to where an eroded surface is generated ^1,3^. This is in contrast to the low OB lineage cell density in the present co-cultures, which at first sight appears a limitation of our model. However, we show that almost all resorption events in the present co-cultures start within the proximity of OBs (since they most likely depend on osteoblastic-RANKL: see above). Thus, our co-culture model and the *in vivo* situations are not that different in this respect, but a low cell density facilitates the visualization of OB visits to resorption events. Of note, *in vivo*, the main osteoblastic partner of OCs is the reversal cell. The present model thus appears an interesting reversal cell model.

As reported, CatK levels greatly vary amongst individuals and depends on factors such as aging, menopause status, or glucocorticoids ^14,22,25^. Intriguingly, these variations correlate inversely with osteoprogenitor recruitment on eroded surfaces and bone formation ^25,36,47^. The present study thus explains this inverse relationship.

### Ectosteric CatK inhibitors to prevent bone loss in the clinic?

From a clinical perspective, it is of interest that the ectosteric inhibitor T06 appears here to inhibit bone resorption and promote osteogenesis as well as ODN, whatever the end point. The advantage of T06 compared to ODN is that it does not affect the homeostasis of growth factors as does ODN^18^ which is believed one of the reasons of ODN’s side effects *in vivo*. Interestingly, T06 was already tested *in vivo*, and shown to increase trabecular structure, osteoblast number, alkaline phosphatase levels and P1NP in ovariectomized mice^18^. Future *in vivo* experiments should further explore osteoclast-osteoblast communication in response to T06, and check whether all ODN’s side effects are avoided.

## Conclusion

It follows from the present study that CatK/collagen degradation should not only be regarded as a determinant of bone resorption – but also as influencing how fast bone formation follows resorption (Figure 6). Several models and clinical trials have stressed that individuals treated with CatK inhibitors have preserved bone formation in contrast with treatments with other resorption antagonists ^27^. Here we provide a rational for this observation and highlight that CatK inhibitors modulate resorptive and osteogenic activity of OCs in opposite directions (Figure 6). These data create awareness of the potential of CatK inhibitors in clinical situations and should encourage the development of ectosteric CatK inhibitors without the side effects of ODN ^18,30,48^.

## Acknowledgements

We thank the members of Dieter Brömme’s lab for helpful discussions in organizing data.

## Author contributions

Preety Panwar; Conceptualization, Data curation, Formal analysis, Investigation, Methodology, Project administration, Software, Supervision, Validation, Visualization, Writing original draft, Writing, review & editing, Funding acquisition; Jacob Olesen: Conceptualization, Data curation, Formal analysis, Methodology, Project administration, Software, Supervision, Writing - review & editing; Jean-Marie Delaisse: Conceptualization, Data curation, Formal analysis, Investigation, Methodology, Supervision, Validation, Visualization, Writing original draft, Writing review & editing; Kent Søe; Conceptualization, Data curation, Formal analysis, Funding acquisition, Investigation, Methodology, Project administration, Resources, Supervision, Visualization, Writing review & editing, Dieter Bromme: Conceptualization, Data curation, Funding acquisition, Investigation, Methodology, Project administration, Resources, Supervision, Validation, Visualization, Writing review & editing.

## Data availability

All data analysed in this study are included in this article and its supplemental materials. Methods in the article contains sufficient information to reproduce reported results.

## Supplemetary material

**Supplementary Figure S1:**
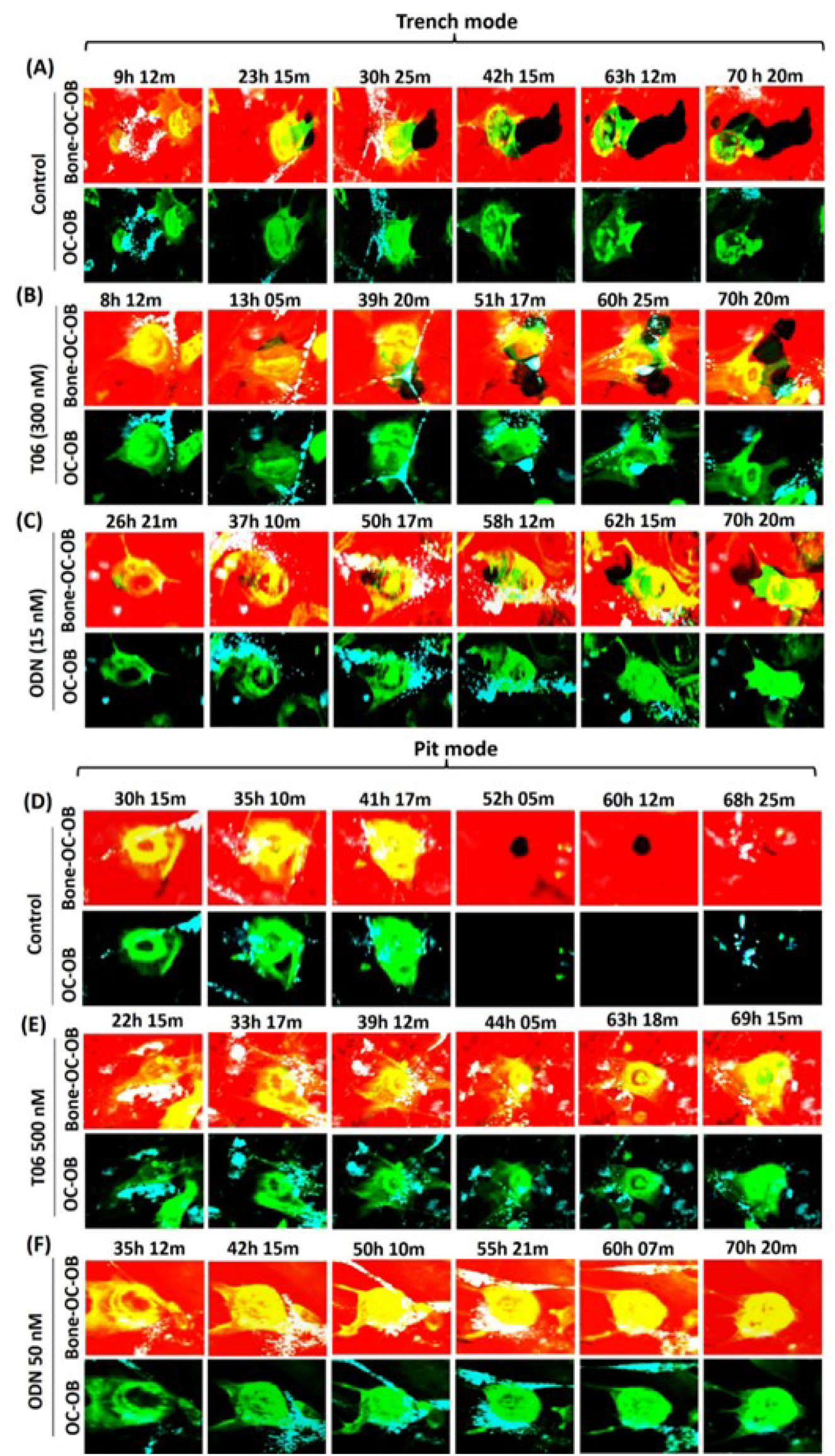
Effect of CatK inhibitors at low concentration on the association of OBs with trenches. (A-C) Selected time-lapse images of OC-OB interaction during trench formation were taken from movies of control (A), T06 300 nM (B), ODN 15 nM (C) (video 2) at various time points over 72 h co-culture. OCs make long trenches in untreated conditions; however, dose dependent inhibition of CatK in OCs showed gradual decrease in size of trenches. OBs showed their preference to excavations generated by CatK-inhibited OCs compared to trenches generated under untreated conditions. (D-E) Effect of CatK inhibitors at high concentration on the association of OBs with pits. Selected time-lapse images of OC-OB interaction during pit formation were taken from videos of control (D), T06 500 nM (E), and ODN 50 nM (F) (video 3) at various time points over 72 h of co-culture. OBs showed their preference to pits generated by OCs subjected to CatK inhibitors (both T06 and ODN) compared to the pits generated under untreated conditions. OCs were stained for actin by using phalloidin (green), OBs were stained with Vybrant DiO cell-labeling solution (cyan) and bone surface was stained with rhodamine (red).

**Supplementary Figure S2.**
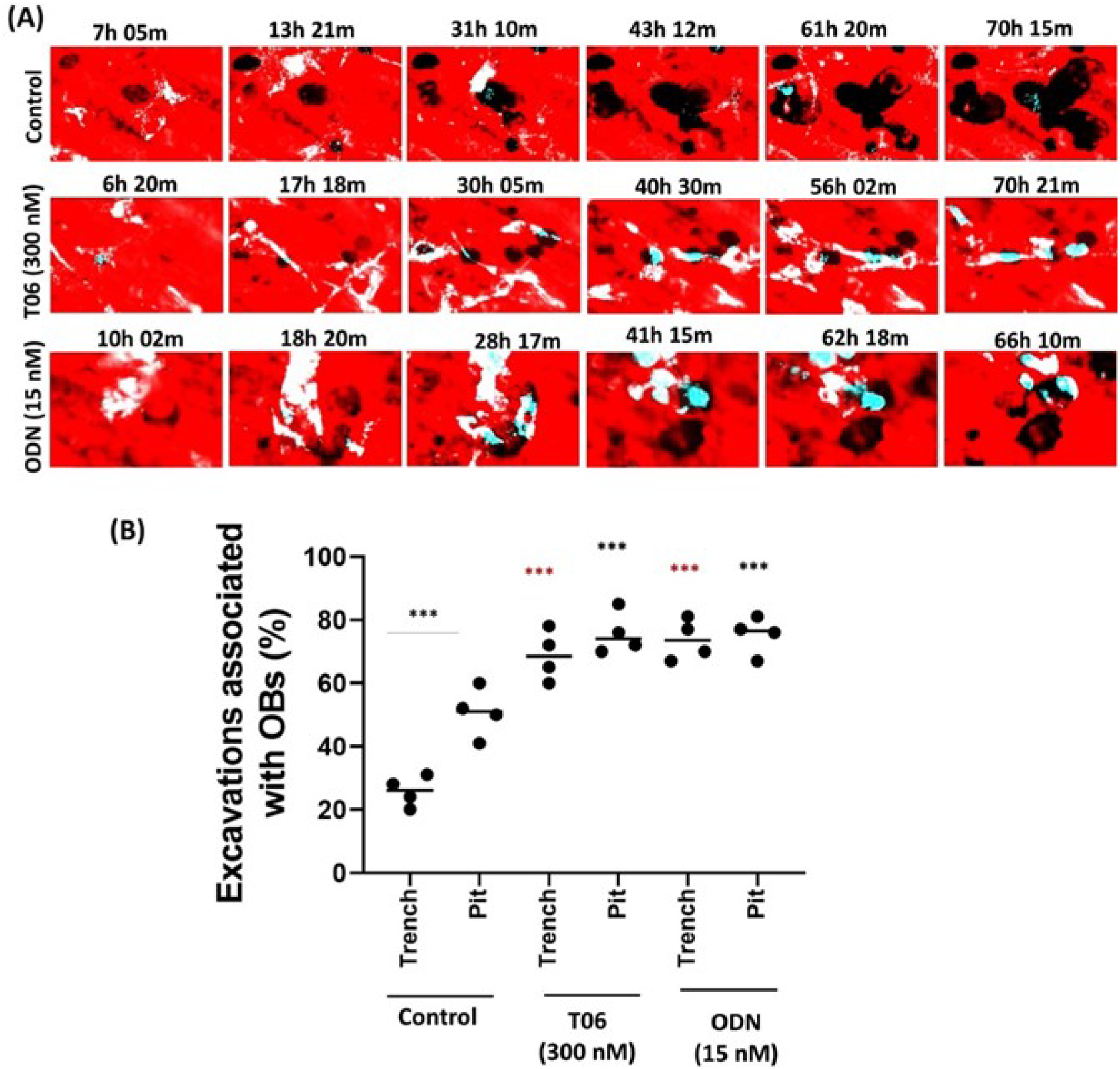
Effect of CatK inhibitors on the association of OBs with resorption events: (A) Time-lapse images showed the interaction of OBs with the excavations in control, T06 (300 nM), and ODN (15 nM) conditions displayed OBs preference towards pits and this interaction increased with CatK inhibition **(supporting data for Fig. 2,3, and 4 and relevant videos; supplementary video 1)**. Sample size: Control: n= 4 donors; 300 nM T06: n=4 donors; 15 nM ODN: n=4 donors). For each donor, 2-3 replicate experiments on individual bone slices were analyzed for all conditions. The median obtained in each experiment are shown as bars. We analyzed between 40-120 OC-OB activities per condition per experiment for each of the donors). (B) Quantification of OB interaction with resorbed cavities under control and inhibitor treated condition. Statistics: Mann-Whitney test (***P < 0.001) to compare the OB interaction with pits or trenches under untreated control condition. Kruskal–Wallis’s test, two tailed (ns: not significant; ***P < 0.0001); Dunn’s multiple comparisons test ***P < 0.001) compared to control. Pits statistics are shown in black and trenches comparison is highlight in red asterisk.

**Supplementary Figure S3:**
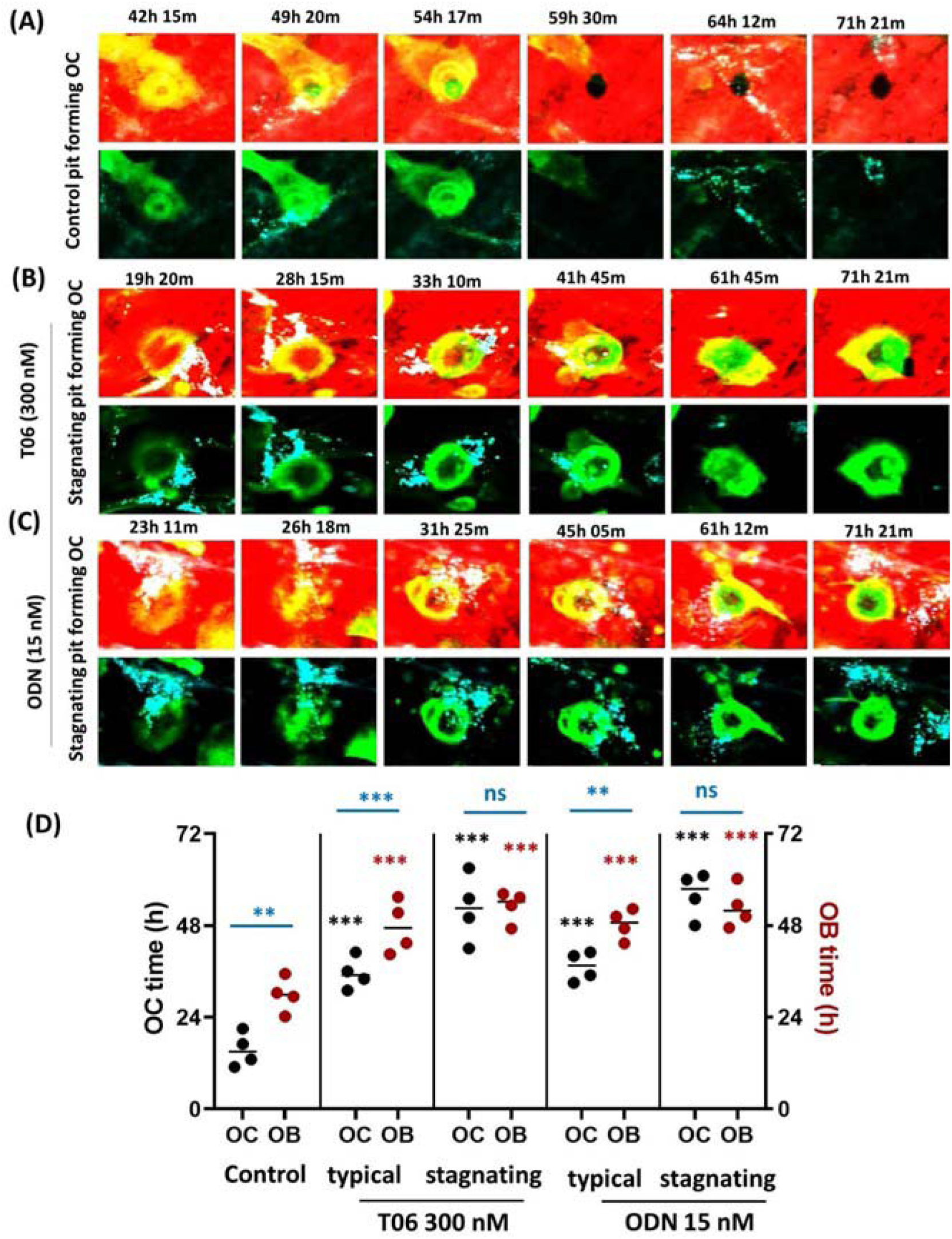
Induction of a stagnating activity state in OCs treated with CatK inhibitors and its consequence for duration of occupancy of the excavation by OCs and OBs. Time-lapse images of cocultures showing the typical resorption behaviour of OCs in generating pit under (A) control, and atypical resorption behavior (B) T06 500nM, and (C) ODN 15 nM treated conditions during a 72 h observation. (D) Duration of presence of OCs and OBs when comparing typical behaviour, stagnating behaviour, and absence of treatment (video 4). Control: n= 4 donors; 300 nM T06: n=4 donors; 15 nM ODN: n=4 donors. For each donor, 2 replicate experiments on individual bone slices were analyzed for all conditions. The median obtained in each experiment are shown as bars. We analyzed between 40-120 OC-OB activities per condition per experiment for each of the donors). Statistics: Kruskal–Wallis’s test, two tailed (ns: not significant; **P < 0.01; ***P < 0.001); Dunn’s multiple comparisons test (ns: not significant; **P < 0.02; ***P < 0.001) compared to the untreated control (OC time: black stars; and OB time red stars). OBs recruitment in pits is also independent of OCs and OBs reside longer duration in pits compared to OCs in both untreated and treated conditions. Statistics: Mann-Whitney test (ns: not significant; **P < 0.01; ***P < 0.001) shown by blue stars.

**Supplementary Figure 4.**
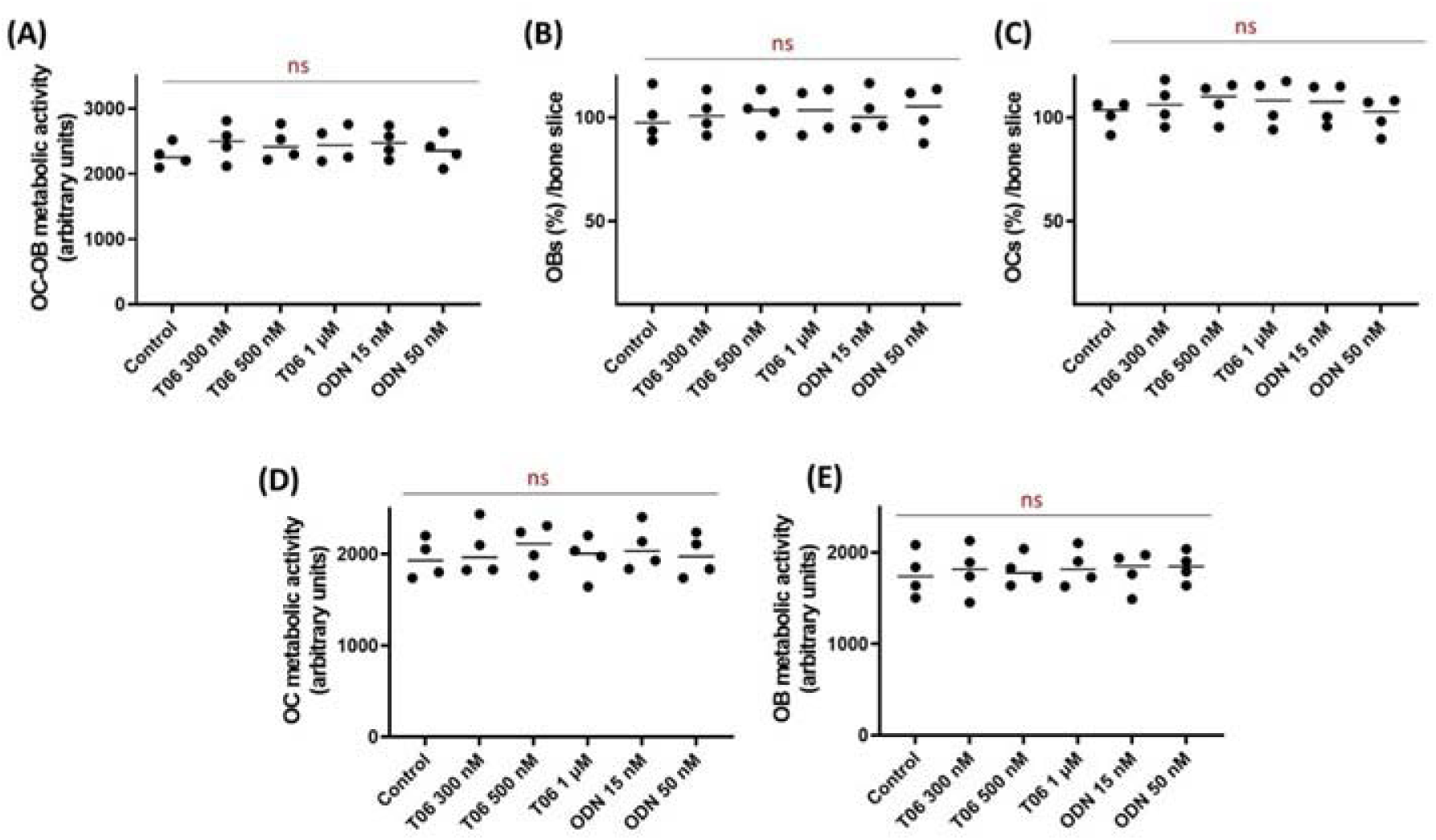
Quantification of (A) metabolic activity, (B) ALP^+^ OBs, and (C)TRACP^+^ OCs in co-culture system grown over bone slice for 72 h in the presence or absence of inhibitors (T06, ODN at different concentrations). Quantification of metabolic activity of (D) OBs, and (E) OCs in mono-culture grown over bone surface to determine the effect of inhibitors (T06, ODN) on the cell viability and proliferation. Sample size (n=4 donors) for control, T06 (300 nM, 500 nM, 1 µM), ODN (15 nM, 50 nM). These data confirm no effect of inhibitors on cells number, viability and proliferation as no significant difference was observed compared to untreated control. Statistics: Kruskal–Wallis’s test, two tailed (ns: not significant) compared with untreated control.

**Supplementary Figure S5:**
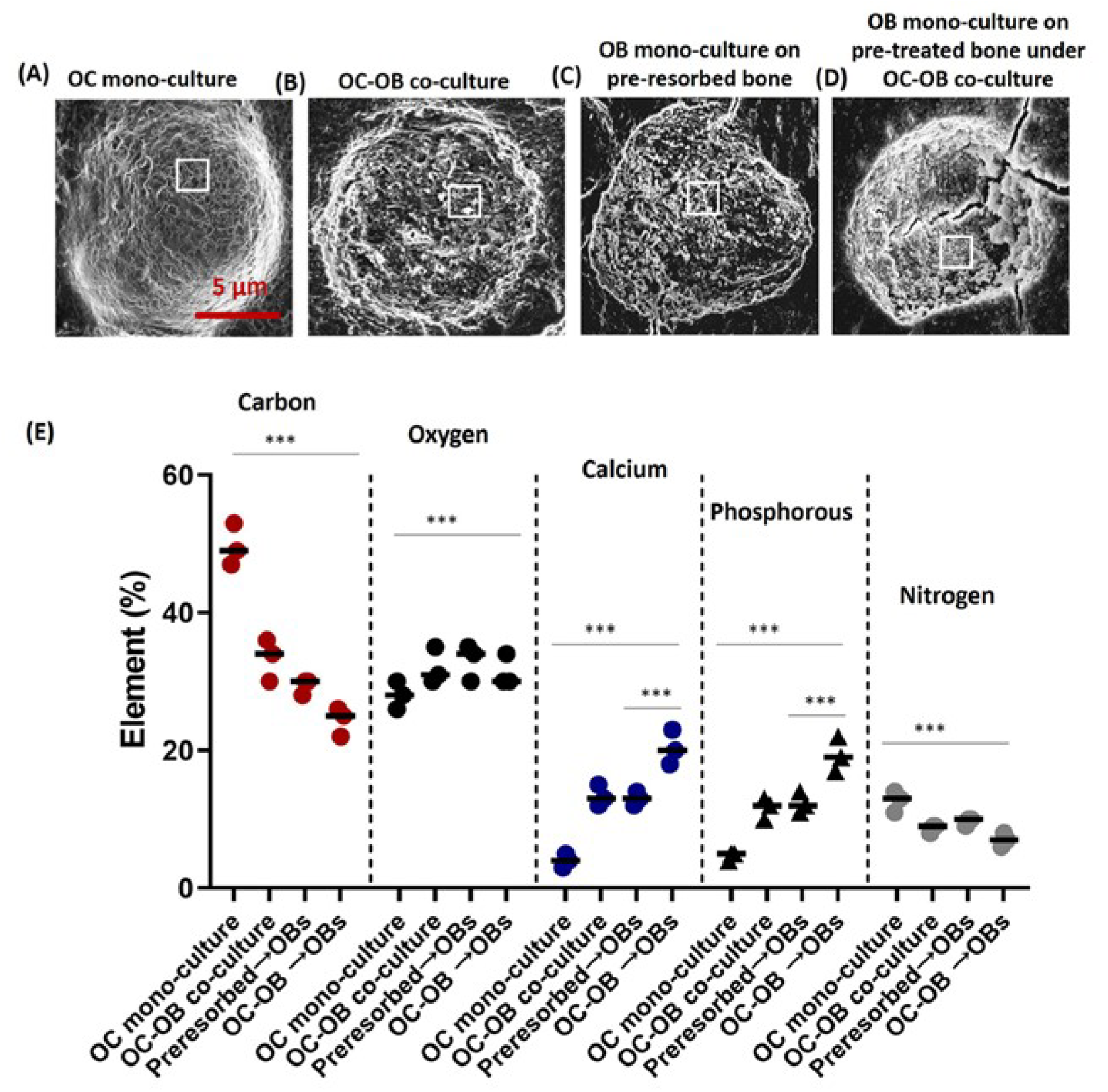
Effect of OBs on the ultrastructure and elemental composition of the resorption pits. (A-C) SEM micrographs (backscattered electron imaging) clearly display the differences in matrix in pits after (A) an OC mono-culture. (B) an OC-OB co-culture, and (C) an OB-culture on pre-resorbed bone slices in co-culture media (D) an OB culture on pre-resorbed bone slice under OC-OB culture in co-culture media. Note the changes in the appearance of the matrix. Scale bar: 5 µm for pits in each condition. (E) Quantification of carbon, oxygen, calcium, phosphorous, and nitrogen in the pits under mono-culture, co-culture, and on pre-resorbed conditions. (5µm^2^ region of interest (ROI); 5-10 regions per pit in each condition for 20 pits). n=3 individual experiments from separate donors. The mean proportions obtained in each experiment are shown as bars. Statistics: Kruskal–Wallis’s test, two tailed (ns: not significant; *P = 0.05, ***P < 0.001); Dunn’s multiple comparisons test (ns: not significant; *P = 0.052, ***P < 0.001) compared with untreated control condition. OC-OB co-culture and OBs cultured on pre-resorbed bone slices showed no significant difference in element accumulation.

**Supplementary Figure S6:**
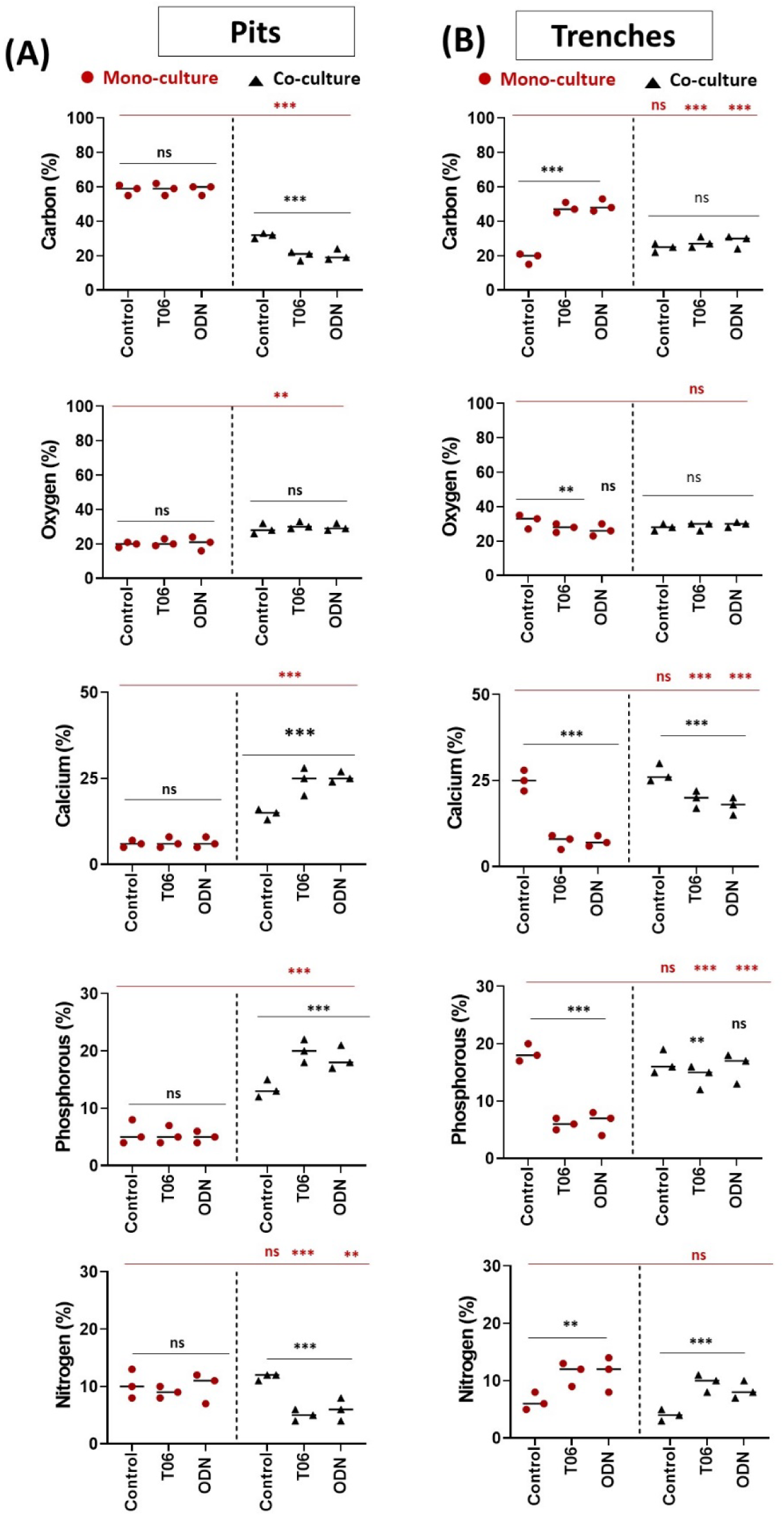
Quantification (atomic percentage) of carbon, oxygen, calcium, phosphorous, and nitrogen in (A) pits and (B) trenches generated in mono-culture and co-cultures with or without CatK inhibitors. The elements were quantified by using energy dispersive X-ray spectroscopy. (Spectrum within 5µm^2^ region of interest (ROI); 5-10 regions per pit in each condition for 25 pits or trenches; n=3 individual experiments from separate donors). The median obtained in each experiment are shown as bars. Statistics: Kruskal–Wallis’s test, two tailed (ns: not significant; **P = 0.01, ***P < 0.001); Dunn’s multiple comparisons test (ns: not significant; **P = 0.008, ***P < 0.001) used to compare the treated conditions with untreated control within mono and co-culture (indicated by black asterisk). Mann–Whitney test was used to compare the elements of mono culture with the co-culture under control and inhibitor treated conditions (indicated by red asterisk). ‘ns’, not significant; **P < 0.01; ***P < 0.001.

**Supplementary Table S1:**
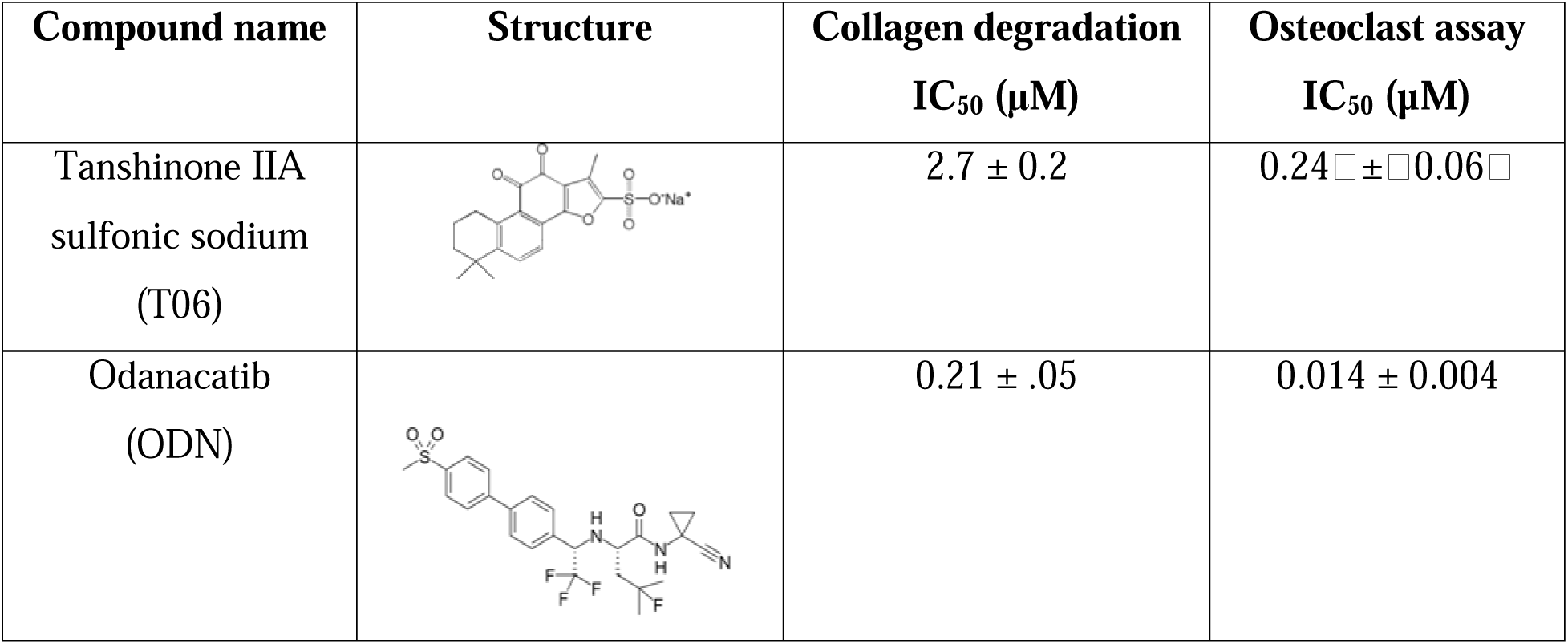
IC_50_ values of ODN and T06 for soluble collagen degradation and bone resorption in osteoclast assay.

| Compound name | Structure | Collagen degradation<br>IC<sub>50</sub> (μM) | Osteoclast assay<br>IC<sub>50</sub> (μM) |
| --- | --- | --- | --- |
| Tanshinone IIA<br>sulfonic sodium<br>(T06) |  | 2.7 ± 0.2 | 0.24 ± 0.06 |
| Odanacatib<br>(ODN) |  | 0.21 ± .05 | 0.014 ± 0.004 |

## REFERENCES

1. Delaisse JM, Andersen TL, Kristensen HB, Jensen PR, Andreasen CM, Søe K. Re-thinking the bone remodeling cycle mechanism and the origin of bone loss. Bone. Dec 2020;141:115628. doi:10.1016/j.bone.2020.115628

2. Abdelgawad ME, Delaisse JM, Hinge M, et al. Early reversal cells in adult human bone remodeling: osteoblastic nature, catabolic functions and interactions with osteoclasts. Histochem Cell Biol. Jun 2016;145(6):603–15. doi:10.1007/s00418-016-1414-y

3. Andersen TL, Jensen PR, Sikjaer TT, Rejnmark L, Ejersted C, Delaisse JM. A Critical Role of the Bone Marrow Envelope in Human Bone Remodeling. J Bone Miner Res. Jun 2023;38(6):918–928. doi:10.1002/jbmr.4815

4. Atkins GJ, Kostakis P, Pan B, et al. RANKL expression is related to the differentiation state of human osteoblasts. J Bone Miner Res. Jun 2003;18(6):1088–98. doi:10.1359/jbmr.2003.18.6.1088

5. El-Masri BM, Andreasen CM, Laursen KS, et al. Mapping RANKL- and OPG-expressing cells in bone tissue: the bone surface cells as activators of osteoclastogenesis and promoters of the denosumab rebound effect. Bone Research. 2024/10/18 2024;12(1):62. doi:10.1038/s41413-024-00362-4

6. Gori F, Hofbauer LC, Dunstan CR, Spelsberg TC, Khosla S, Riggs BL. The expression of osteoprotegerin and RANK ligand and the support of osteoclast formation by stromal-osteoblast lineage cells is developmentally regulated. Endocrinology. Dec 2000;141(12):4768–76. doi:10.1210/endo.141.12.7840

7. Matic I, Matthews BG, Wang X, et al. Quiescent Bone Lining Cells Are a Major Source of Osteoblasts During Adulthood. Stem Cells. Dec 2016;34(12):2930–2942. doi:10.1002/stem.2474

8. Pirapaharan DC, Olesen JB, Andersen TL, et al. Catabolic activity of osteoblast lineage cells contributes to osteoclastic bone resorption in vitro. J Cell Sci. May 15 2019;132(10)doi:10.1242/jcs.229351

9. Streicher C, Heyny A, Andrukhova O, et al. Estrogen Regulates Bone Turnover by Targeting RANKL Expression in Bone Lining Cells. Sci Rep. Jul 25 2017;7(1):6460. doi:10.1038/s41598-017-06614-0

10. Sims NA, Martin TJ. Osteoclasts Provide Coupling Signals to Osteoblast Lineage Cells Through Multiple Mechanisms. Annu Rev Physiol. Feb 10 2020;82:507–529. doi:10.1146/annurev-physiol-021119-034425

11. Teti A. Mechanisms of osteoclast-dependent bone formation. Bonekey Rep. Dec 4 2013;2:449. doi:10.1038/bonekey.2013.183

12. Owen TA, Aronow M, Shalhoub V, et al. Progressive development of the rat osteoblast phenotype in vitro: reciprocal relationships in expression of genes associated with osteoblast proliferation and differentiation during formation of the bone extracellular matrix. J Cell Physiol. Jun 1990;143(3):420–30. doi:10.1002/jcp.1041430304

13. Everts V, Delaissé JM, Korper W, et al. The bone lining cell: its role in cleaning Howship’s lacunae and initiating bone formation. J Bone Miner Res. Jan 2002;17(1):77–90. doi:10.1359/jbmr.2002.17.1.77

14. Borggaard XG, Pirapaharan DC, Delaissé J-M, Søe K. Osteoclasts’ Ability to Generate Trenches Rather Than Pits Depends on High Levels of Active Cathepsin K and Efficient Clearance of Resorption Products. International Journal of Molecular Sciences. 2020;21(16):5924.

15. Delaisse JM, Søe K, Andersen TL, Rojek AM, Marcussen N. The Mechanism Switching the Osteoclast From Short to Long Duration Bone Resorption. Front Cell Dev Biol. 2021;9:644503. doi:10.3389/fcell.2021.644503

16. Panwar P, Olesen JB, Blum G, Delaisse J-M, Søe K, Brömme D. Real-time analysis of osteoclast resorption and fusion dynamics in response to bone resorption inhibitors. Scientific Reports. 2024/03/28 2024;14(1):7358. doi:10.1038/s41598-024-57526-9

17. Panwar P, Søe K, Guido RV, Bueno RVC, Delaisse J-M, Brömme D. A novel approach to inhibit bone resorption: exosite inhibitors against cathepsin K. British Journal of Pharmacology. 2016;173(2):396–410. doi:10.1111/bph.13383

18. Panwar P, Xue L, Søe K, et al. An Ectosteric Inhibitor of Cathepsin K Inhibits Bone Resorption in Ovariectomized Mice. J Bone Miner Res. Dec 2017;32(12):2415–2430. doi:10.1002/jbmr.3227

19. Søe K, Delaissé JM. Time-lapse reveals that osteoclasts can move across the bone surface while resorbing. J Cell Sci. Jun 15 2017;130(12):2026–2035. doi:10.1242/jcs.202036

20. Søe K, Merrild DM, Delaissé JM. Steering the osteoclast through the demineralization- collagenolysis balance. Bone. Sep 2013;56(1):191–8. doi:10.1016/j.bone.2013.06.007

21. Gentzsch C, Delling G, Kaiser E. Microstructural classification of resorption lacunae and perforations in human proximal femora. Calcif Tissue Int. Jun 2003;72(6):698–709. doi:10.1007/s00223-002-2020-7

22. Merrild DM, Pirapaharan DC, Andreasen CM, et al. Pit- and trench-forming osteoclasts: a distinction that matters. Bone Res. 2015;3:15032. doi:10.1038/boneres.2015.32

23. Mulari M, Vaaraniemi J, Vaananen HK. Intracellular membrane trafficking in bone resorbing osteoclasts. Microsc Res Tech. Aug 15 2003;61(6):496–503. doi:10.1002/jemt.10371

24. Rumpler M, Würger T, Roschger P, et al. Osteoclasts on bone and dentin in vitro: mechanism of trail formation and comparison of resorption behavior. Calcif Tissue Int. Dec 2013;93(6):526–39. doi:10.1007/s00223-013-9786-7

25. Møller AMJ, Delaissé J-M, Olesen JB, et al. Aging and menopause reprogram osteoclast precursors for aggressive bone resorption. Bone Research. 2020a;8(1)doi:10.1038/s41413-020-0102-7

26. Søe K, Delaissé J-M. Glucocorticoids maintain human osteoclasts in the active mode of their resorption cycle. Journal of Bone and Mineral Research. 2010;25(10):2184–2192. doi:10.1002/jbmr.113

27. Drake MT, Clarke BL, Oursler MJ, Khosla S. Cathepsin K Inhibitors for Osteoporosis: Biology, Potential Clinical Utility, and Lessons Learned. Endocr Rev. Aug 1 2017;38(4):325–350. doi:10.1210/er.2015-1114

28. Rünger TM, Adami S, Benhamou C-L, et al. Morphea-like skin reactions in patients treated with the cathepsin K inhibitor balicatib. Journal of the American Academy of Dermatology. 2012/03/01/ 2012;66(3):e89-e96. 10.1016/j.jaad.2010.11.033

29. Frangogiannis N. Transforming growth factor-β in tissue fibrosis. J Exp Med. Mar 2 2020;217(3):e20190103. doi:10.1084/jem.20190103

30. Panwar P, Law S, Jamroz A, et al. Tanshinones that selectively block the collagenase activity of cathepsin K provide a novel class of ectosteric antiresorptive agents for bone. Br J Pharmacol. Mar 2018;175(6):902–923. doi:10.1111/bph.14133

31. Møller AMJ, Delaissé J-M, Søe K. Osteoclast Fusion: Time-Lapse Reveals Involvement of CD47 and Syncytin-1 at Different Stages of Nuclearity. Journal of Cellular Physiology. 2017;232(6):1396–1403. doi:10.1002/jcp.25633

32. Delaisse JM. The reversal phase of the bone-remodeling cycle: cellular prerequisites for coupling resorption and formation. Bonekey Rep. 2014;3:561. doi:10.1038/bonekey.2014.56

33. Leung P, Pickarski M, Zhuo Y, Masarachia PJ, Duong LT. The effects of the cathepsin K inhibitor odanacatib on osteoclastic bone resorption and vesicular trafficking. Bone. Oct 2011;49(4):623–35. doi:10.1016/j.bone.2011.06.014

34. Abdelgawad ME, Søe K, Andersen TL, et al. Does collagen trigger the recruitment of osteoblasts into vacated bone resorption lacunae during bone remodeling? Bone. Oct 2014;67:181–8. doi:10.1016/j.bone.2014.07.012

35. Lassen NE, Andersen TL, Pløen GG, et al. Coupling of Bone Resorption and Formation in Real Time: New Knowledge Gained From Human Haversian BMUs. J Bone Miner Res. Jul 2017;32(7):1395–1405. doi:10.1002/jbmr.3091

36. Jensen PR, Andersen TL, Pennypacker BL, Duong LT, Delaissé JM. The bone resorption inhibitors odanacatib and alendronate affect post-osteoclastic events differently in ovariectomized rabbits. Calcif Tissue Int. Feb 2014;94(2):212–22. doi:10.1007/s00223-013-9800-0

37. Fuller K, Lawrence KM, Ross JL, et al. Cathepsin K inhibitors prevent matrix-derived growth factor degradation by human osteoclasts. Bone. Jan 2008;42(1):200–11. doi:10.1016/j.bone.2007.09.044

38. Lotinun S, Kiviranta R, Matsubara T, et al. Osteoclast-specific cathepsin K deletion stimulates S1P-dependent bone formation. J Clin Invest. Feb 2013;123(2):666–81. doi:10.1172/jci64840

39. Xie H, Cui Z, Wang L, et al. PDGF-BB secreted by preosteoclasts induces angiogenesis during coupling with osteogenesis. Nat Med. Nov 2014;20(11):1270–8. doi:10.1038/nm.3668

40. Parikka V, Peng Z, Hentunen T, et al. Estrogen responsiveness of bone formation in vitro and altered bone phenotype in aged estrogen receptor-α-deficient male and female mice. European Journal of Endocrinology. 2005;152(2):301–314. doi:10.1530/eje.1.01832

41. Mulari MT, Qu Q, Härkönen PL, Väänänen HK. Osteoblast-like cells complete osteoclastic bone resorption and form new mineralized bone matrix in vitro. Calcif Tissue Int. Sep 2004;75(3):253–61. doi:10.1007/s00223-004-0172-3

42. Kiviranta R, Morko J, Alatalo SL, et al. Impaired bone resorption in cathepsin K-deficient mice is partially compensated for by enhanced osteoclastogenesis and increased expression of other proteases via an increased RANKL/OPG ratio. Bone. Jan 2005;36(1):159–72. doi:10.1016/j.bone.2004.09.020

43. Andersen TL, del Carmen Ovejero M, Kirkegaard T, Lenhard T, Foged NT, Delaissé JM. A scrutiny of matrix metalloproteinases in osteoclasts: evidence for heterogeneity and for the presence of MMPs synthesized by other cells. Bone. Nov 2004;35(5):1107-19. doi:10.1016/j.bone.2004.06.019

44. Fuller K, Chambers TJ. Localisation of mRNA for collagenase in osteocytic, bone surface and chondrocytic cells but not osteoclasts. J Cell Sci. Jun 1995;108 ( Pt 6):2221–30. doi:10.1242/jcs.108.6.2221

45. Delaissé JM, Eeckhout Y, Neff L, et al. (Pro)collagenase (matrix metalloproteinase-1) is present in rodent osteoclasts and in the underlying bone-resorbing compartment. J Cell Sci. Dec 1993;106 ( Pt 4):1071–82. doi:10.1242/jcs.106.4.1071

46. Garnero P, Ferreras M, Karsdal MA, et al. The type I collagen fragments ICTP and CTX reveal distinct enzymatic pathways of bone collagen degradation. J Bone Miner Res. May 2003;18(5):859–67. doi:10.1359/jbmr.2003.18.5.859

47. Andersen TL, Abdelgawad ME, Kristensen HB, et al. Understanding coupling between bone resorption and formation: are reversal cells the missing link? The American journal of pathology. 2013;183(1):235–246.

48. Brömme D, Panwar P, Turan S. Cathepsin K osteoporosis trials, pycnodysostosis and mouse deficiency models: Commonalities and differences. Expert Opin Drug Discov. 2016;11(5):457–72. doi:10.1517/17460441.2016.1160884

